# Distinct contributions of motor imagery and execution to history-dependent biases in reaching

**DOI:** 10.64898/2026.04.17.719269

**Authors:** Christian Seegelke, Tobias Heed

**Author notes:** Correspondence Dr. Christian Seegelke University of Salzburg Department of Psychology Hellbrunner Straße 34, 5020 Salzburg, Austria.

## Abstract

The recent movement history shapes motor performance, that is, previous movements can affect current movement characteristics such as trajectory shape. History effects are commonly attributed to carryover of motor-related activity. However, action execution entails sensory feedback; therefore, an alternative is that history effects stem from the sensory information produced by previous movements. To dissociate motor and sensory contributions, we assessed whether history effects emerge from imagined movements, which involve movement planning but not sensory feedback. Overt reaches around an obstacle led to systematic adjustment of the initial reach direction of following reaches – a hallmark of motor history. Imagined reaches around obstacles induced similar biases, albeit with smaller magnitude, presumably due to the need to inhibit overt execution during imagery. By contrast, execution but not imagery induced biases in late, feedback-related measures, suggesting that these history effects depended on sensory rather than motor aspects of the movement history. Thus, motor and sensory signals make distinct and complementary contributions to movement history: recent motor states shape feedforward planning, whereas recent sensory states shape feedback-related movement refinement.

## Introduction

Motor performance is shaped not only by the current state of the body and environment but also by movements that occurred in the recent past. Such history-dependent effects have been observed across a wide range of tasks and manifest as systematic biases in movement parameters, including grasp orientation (Schütz et al., 2011; Seegelke et al., 2015, 2013), reach direction (Diedrichsen et al., 2010; Tsay et al., 2022; Verstynen and Sabes, 2011), movement speed (Hammerbeck et al., 2014), and response latency (Mawase et al., 2018; Wong et al., 2017). Notably, these effects can arise even after only a single preceding trial (Jax and Rosenbaum, 2007; Seegelke et al., 2021; Seegelke and Heed, 2025; Valyear and Frey, 2014). For example, a center-out reach was more curved if participants had reached around an obstacle in the previous movement than if the previous reach had been unobstructed (Jax and Rosenbaum, 2009, 2007).

History-dependent biases are commonly attributed to carryover of motor-related activity from prior movements (Jax and Rosenbaum, 2007; Rosenbaum et al., 2007). However, such biases can also arise when the preceding movement is merely observed rather than performed (Griffiths and Tipper, 2012, 2009; Seegelke et al., 2013). Because action observation and movement execution recruit overlapping premotor and parietal regions (Hardwick et al., 2018), observation-induced biases might reflect motor simulation through activation of the observer’s mirror neuron system (Rizzolatti and Craighero, 2004). Yet, since both movement execution and action observation involve sensory feedback, an alternative explanation is that history effects instead arise from the sensory information associated with the executed or observed movement, rather than from motor simulation per se.

Motor Imagery, that is, the mental rehearsal of a movement without physically performing it (Decety and Grèzes, 1999), provides a way to dissociate between these two alternatives. Behavioral, physiological, and neuroimaging studies have supported the view that executed and imagined movements are functionally equivalent (Jeannerod, 1994; Jeannerod and Decety, 1995): they exhibit similarities in timing (Decety et al., 1989; Sirigu et al., 1996), speed-accuracy tradeoff (Jeannerod and Decety, 1995; Sirigu et al., 1996), and physiological responses (Decety et al., 1993, 1991), and they share activation in common (pre-)motor, parietal, and cerebellar areas (Hardwick et al., 2018; Hétu et al., 2013; Lotze and Halsband, 2006). Moreover, TMS-induced modulations of corticospinal excitability are muscle-specific and task-dependent after both executed and imagined movements (Fadiga et al., 1998; Grosprêtre et al., 2016; Kasai et al., 1997; Traverse et al., 2018). In addition, motor imagery can support motor learning and improve motor performance (Anwar et al., 2011; Gippert et al., 2025; Ladda et al., 2021; Sheahan et al., 2018) and can induce cortical plasticity akin to overt execution (Avanzino et al., 2015; Ruffino et al., 2019, 2017). Finally, motor imagery produces the same sensory attenuation effects as real movements, indicating that the brain employs sensory predictions even without overt execution. This suggests that motor imagery engages feedforward motor commands in a way that closely resembles overt execution (Kilteni et al., 2018). These multiple parallels between the processing of overt and imagined movements are typically interpreted as the two sharing their underlying mechanisms.

The crucial difference between executed and imagined movement is that the latter produces no movement-related sensory feedback. Consequently, any difference in history effects between execution and imagery could be attributed to the contribution of sensory feedback from the prior movement. Following this logic, we tested whether reach trajectories exhibit comparable biases after a preceding reach around an obstacle, irrespective of whether that prior movement was performed overtly or only imagined. History effects have typically been examined only with respect to feedforward-related movement aspects – that is, early movement characteristics, such as the reach’s initial direction, that cannot yet have been modified based on sensory feedback. Here, we additionally assessed whether history effects also emerge for movement aspects later in the reach that are susceptible to online correction based on sensory feedback.

We observed that motor and sensory aspects of the prior movement made distinct contributions to the emergent motor history. Early, feedforward-related movement characteristics were affected by a motor history established both by execution and imagery, suggesting they reflect a motor-based history; in contrast, late, feedback-related movement characteristics were sensitive only to the history of executed movements, thus suggesting a sensory-based history. Accordingly, motor and sensory signals make distinct contributions in shaping subsequent motor performance.

## Results

In Experiment 1, participants performed two consecutive center-out reaching movements, referred to as prime and probe movement, to move the cursor of a mouse from a start position to a target position (execution trials). On some trials an obstacle was placed halfway between the start and target position, and to avoid it, participants had to produce curved trajectories (Fig. 1A, Execution). On a subset of trials (motor imagery trials) participants were cued to imagine the prime movement but withhold its execution (Fig. 1A, Motor Imagery). We measured participants’ absolute initial reach error in probe movements to determine how strongly movements deviated from a straight path at reach start. During this early part of the movement, feedback – be it visual or proprioceptive – cannot yet have reached the brain, so that modulation of initial reach angle must stem from feedforward, that is, motor planning-related processing. We quantified history effects as the difference in initial reach error between probe movements for which the prime movement had, vs. had not, involved an obstacle; we refer to this measure as initial reach bias.

**Figure 1.**
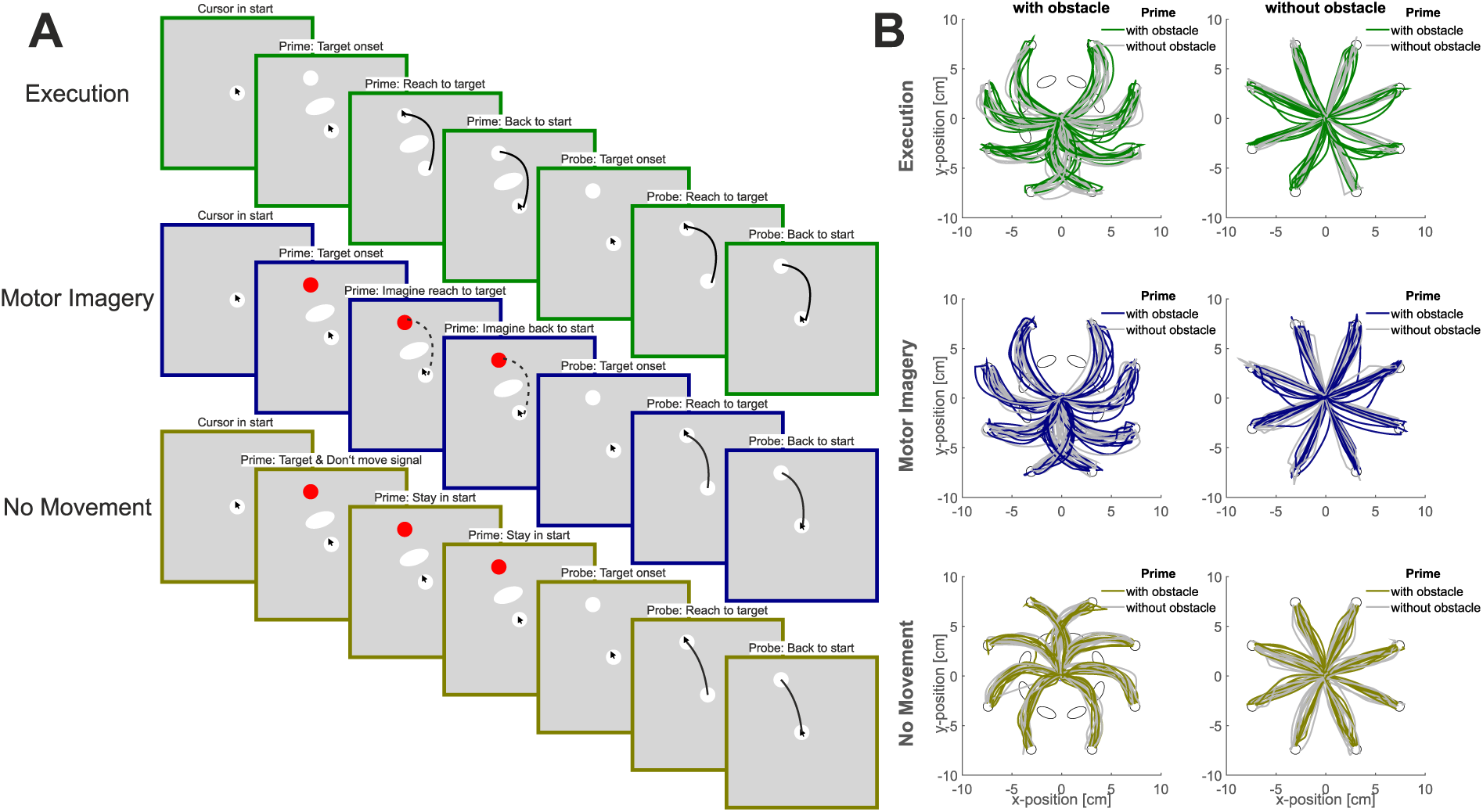
Trial structure and reach trajectories. **A** Exemplary trial structure of an Execution trial (top row), a Motor Imagery trial (middle row), and a No Movement trial (bottom row). The plot shows trials in which an obstacle was present during the prime but not during the probe phase. **B** Reach trajectories of an exemplary participant for all target-obstacle combinations. Only a single target and obstacle were present in any given trial (see A). Probe movements exhibited larger curvature when the previous (i.e., prime) movement contained an obstacle (colored traces) than when it did not (grey traces).

### Feedforward control is shaped by prior motor imagery

Replicating previous work (Jax and Rosenbaum, 2007; Seegelke and Heed, 2026), participants exhibited a strong reach bias in probe movements, evidenced in larger initial reach errors when the preceding, executed movement (i.e., the prime movement) required clearing an obstacle (Fig. 1B, 2AB green data). This was true regardless of whether the probe movement required obstacle avoidance (probe with obstacle: reach bias = 6.7°, [5.0,8.3], brackets contain 95% highest density interval, pd > 99%, 0% in ROPE) or not (probe without obstacle: reach bias = 12.8°, [10.0,15.7], pd > 99%, 0% in ROPE). The bias was present in every individual (Fig. 2EF green data).

**Figure 2.**
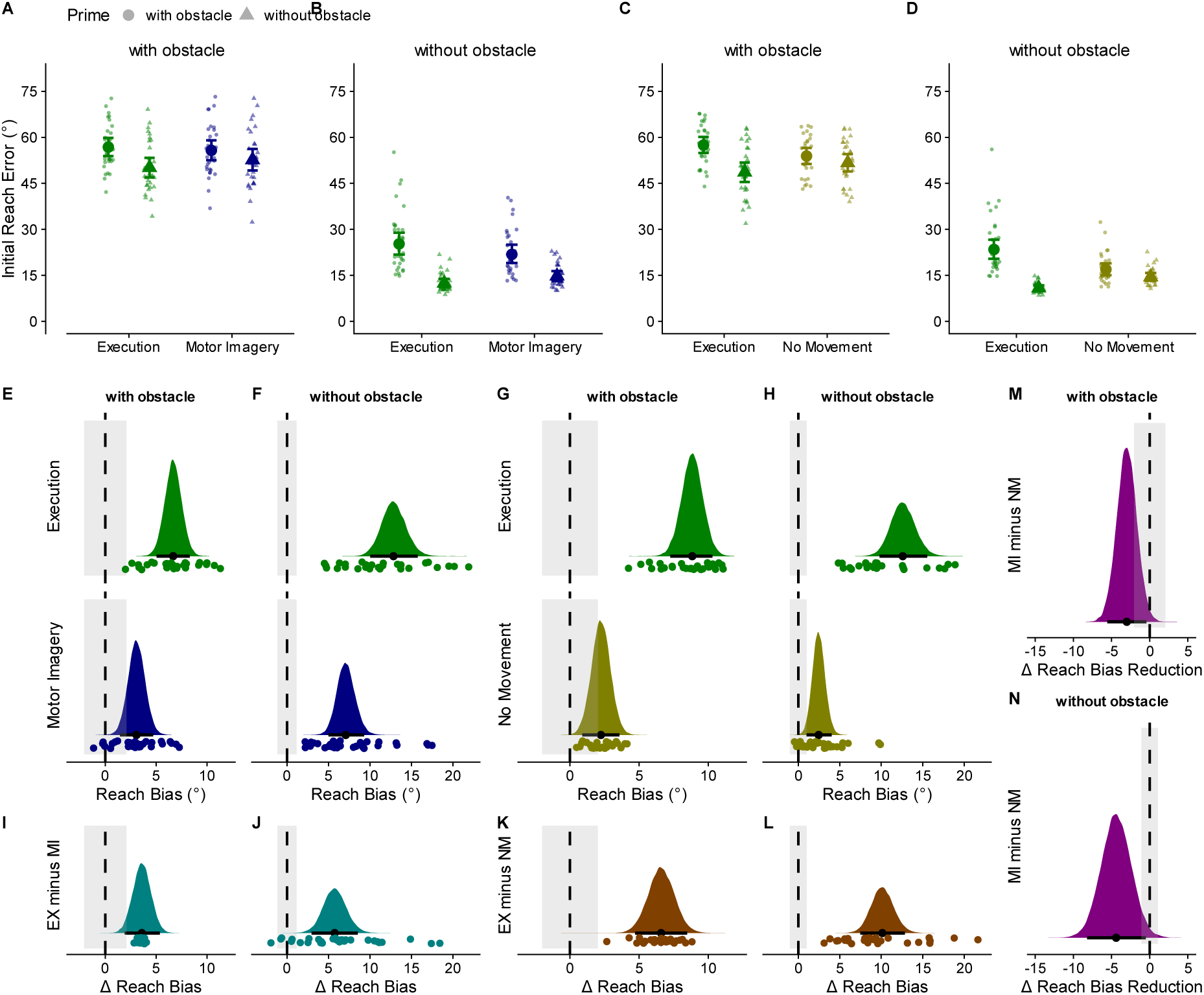
History effects of prime movement on feedforward-related probe movement aspects in Experiment 1 (Motor Imagery) and Experiment 2 (No Movement). **A-D** Initial reach error for Experiment 1 (AB) and Experiment 2 (CD). Large symbols represent the group estimates and error bars the 95% highest-density interval (HDI). Small symbols represent individual participant estimates. **E-H** Reach bias, calculated as difference of initial reach error in trials with minus without an obstacle in the prime movement for Execution and Motor Imagery (EF) and No Movement (GH) for probe movements with and without an obstacle. **I-L** Difference in reach bias between Execution [EX] and Motor Imagery [MI] (IJ) and No Movement [NM] trials (KL), respectively. **M-N** Difference in reach bias reduction between Experiment 1 (Motor Imagery) and Experiment 2 (No Movement). Colored areas: posterior distributions. Black dots: median; error bars: 95% highest-density interval (HDI); grey shaded areas: region of practical equivalence (ROPE).

Importantly, when participants only imagined the prime movement, the bias was also present (probe with obstacle: reach bias = 3.1°, [1.4,4.7], pd > 99%, 19% in ROPE; probe without obstacle: reach bias = 7.1°, [5.0,9.3], pd > 99%, 0% in ROPE; 2ABEF blue data), though smaller in size (probe with obstacle: difference in reach bias = 3.6°, [1.9,5.4], pd > 99%, 5% in ROPE; probe without obstacle: difference in reach bias = 5.7°, [3.0,8.5], pd > 99%, 0% in ROPE; Fig. 2IJ). Thus, motor imagery is sufficient to bias feedforward control of subsequent movements.

### Motor imagery induced reach bias is not due to visuomotor priming

To corroborate that the observed reach bias is attributable to motor imagery and not to visuomotor priming elicited by the repeated visual presentation of the obstacle (Hesse et al., 2008; Seegelke et al., 2016), we contrasted our results with a control condition (Experiment 2), in which participants saw, in a subset of trials, the target and obstacle just like in other trials, but target color instructed them to refrain from planning and executing the prime movement (Fig. 1AB, yellow data). This condition, thus, creates a repetition of the visual context, or a visual history, that matches the visual context of all other conditions. As in our imagery condition, no overt movement occurred, but in addition – and in contrast to imagery – the movement was not imagined.

A small bias was present in the No Movement condition (probe with obstacle: reach bias = 2.2°, [0.9,3.6], pd > 99%, 42% in ROPE; probe without obstacle: reach bias = 2.5°, [1.0,4.0], pd > 99%, 1% in ROPE; Fig. 2CDGH) but the difference in reach bias from execution, that is, reach bias after Execution primes minus reach bias in Motor Imagery/ No Movement primes was smaller for Motor Imagery than for No Movement (probe with obstacle: difference in reach bias reduction = -3.0°, [-5.6,-0.4], pd = 99%, 32% in ROPE; without obstacle: difference in reach bias reduction = -4.4°, [-8.2,-0.5], pd = 99%, 7% in ROPE; Fig. 2MN). This comparison indicates that a minor portion of the history bias, namely the portion evident in the No Movement condition, emerges from visuomotor priming; the larger part of the history bias observed after imagined movements, however, can be attributed to the imagined execution of the movement.

### Chronometric properties of prime movements suggest common feedforward mechanisms

The size difference of the history effect in execution vs. imagery may reflect that the two conditions involve procedural, qualitative differences. Alternatively, imagined prime movements may have been less similar to probe movements than executed prime movements, thus reducing the effect of their history on probes. To assess this latter possibility, we compared the duration of executed and imagined prime movements (Decety et al., 1989; Sirigu et al., 1996). Prime duration was longer for execution than imagery prime movements (median difference = 405 ms, [251,542], pd > 99%, 0% in ROPE), and imagery exhibited considerably more variability (Fig. 3AC). Yet, prime duration was longer for movements with than without an obstacle for both execution and imagery (execution: median difference = 95 ms, [71,119], pd > 99%, 0% in ROPE; motor imagery: median difference = 81 ms, [54,109], pd > 99%, 0% in ROPE; Fig. 3B). The movement time difference was, moreover, comparable in the two conditions (median difference in prime duration differences = 14 ms, [-12,40], pd = 85%, 100% in ROPE; Fig. 3D) and moderately correlated (r = 0.48, [0.18,0.69], pd > 99%; Fig. 3E). Overall, this result pattern is consistent with the conclusion that feedforward-related motor history emerges from a common process during movement execution and imagery which suffers from greater variability for imagined than executed movements, which limits the expression of history effect on subsequent movements.

**Figure 3.**
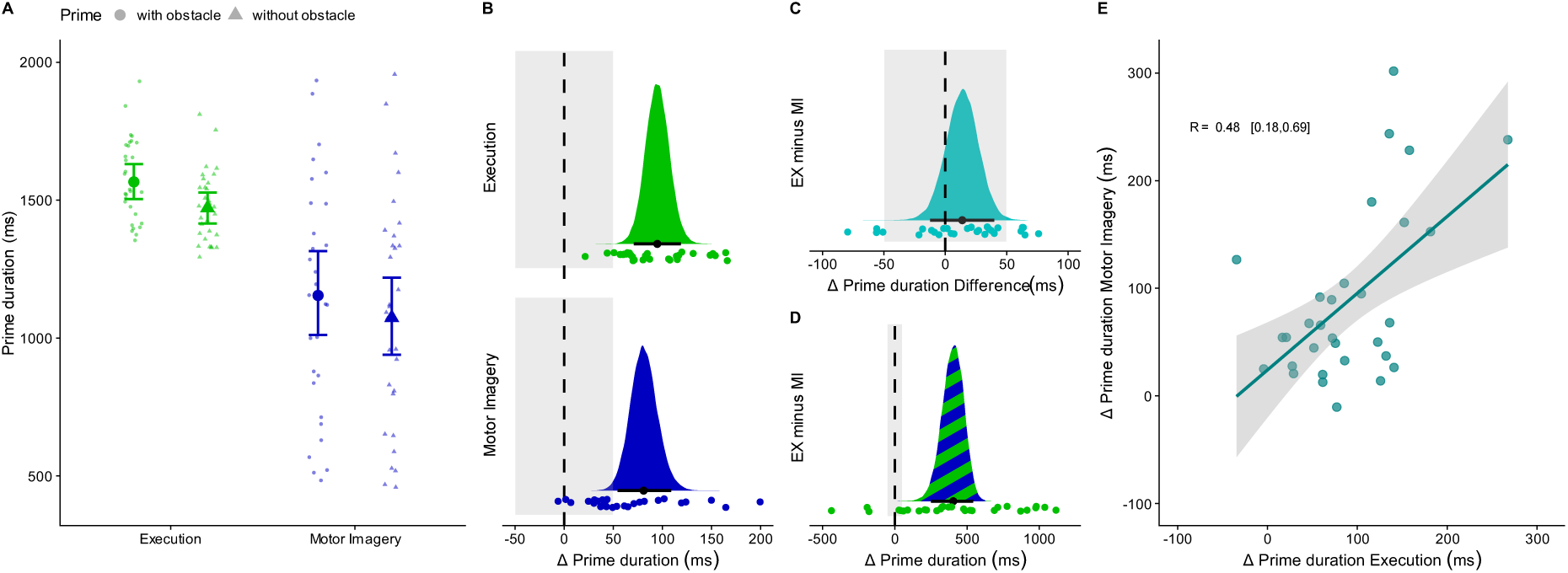
Prime movement times in Experiment 1. **A** Prime movement duration as a function of movement type (Execution, Motor Imagery) and prime context (with vs. without obstacle). **B-D** Contrasts. Overall, prime duration was longer for Execution than Motor Imagery (C), but for both movement types, prime duration was longer with than without an obstacle (B), and this difference was of similar size (D). **E** Correlation between difference in prime duration for movements with minus without an obstacle for Execution and Motor Imagery. Teal line: linear fit; grey shading: ±1 s.e. of the fit. All other details are as in Figure 2.

We further assessed whether the behavioral effects of imagery-induced motor history relate to the self-reported tendency and ability to use motor imagery. However, we observed no reliable correlations between the reach bias and the Motor Imagery Questionnaire (MIQ-RS) scores (r between 0.01 and 0.29, all pd < 97%), the ease of motor imagery (r between -0.07 and 0.14, all pd < 97%), or the frequency of motor imagery (r between -0.23 and 0.03, all pd <97%; see Table S1 and Fig. S1).

### Expression of history effects in feedback control requires overt motor execution

In our characterization of history effects, we have so far focused on feedforward-related movement aspects. Based on the hypothesis that feedforward and feedback processes are governed by a common neural circuitry (Pruszynski, 2014; Scott, 2016), we extended our current experimental approach to investigate whether the movement history also affects later movement phases that rely on sensory feedback. We reasoned that if history effects manifest in feedback-related aspects, movement performance should be improved when the current movement is similar to the preceding one – that is, both prime and probe involve an obstacle, or both prime and probe do not involve an obstacle. We operationalized feedback-related movement aspects as the deviation of probe movements from a straight trajectory at the end of the movement, quantified as the absolute angular deviation between a line connecting cursor position at reach onset and target center and a line connecting cursor position at reach onset and cursor position at target hit. This quantification is analogous to the feedforward-related, initial reach error, and we refer to it as final reach error.

Final reach error was smaller when prime and probe required similar movements (probe with obstacle: difference = -0.06°, [-0.10,-0.02], pd > 99%, 91% in ROPE; probe without obstacle: difference = 0.13°, [0.07,0.19], pd > 99%, 25% in ROPE; Fig. 4EF green data). There was no statistical difference in imagery trials (probe with obstacle: difference = 0.04°, [-0.00,0.09], pd < 97%, 100% in ROPE; probe without obstacle: difference = 0.05°, [-0.02,0.11], pd = 92%, 88% in ROPE; Fig. 4EF blue data). Direct comparison of execution and imagery trials confirmed a stronger influence of the overtly executed than imagined prime movements, though only in trials with an obstacle (difference in final error difference = -0.10°, [-0.16,-0.04], pd > 99%, 44% in ROPE), but not in trials without an obstacle (difference in final error difference = 0.08°, [-0.01,0.17], pd = 96%, 59% in ROPE; Fig. 4IJ).

**Figure 4.**
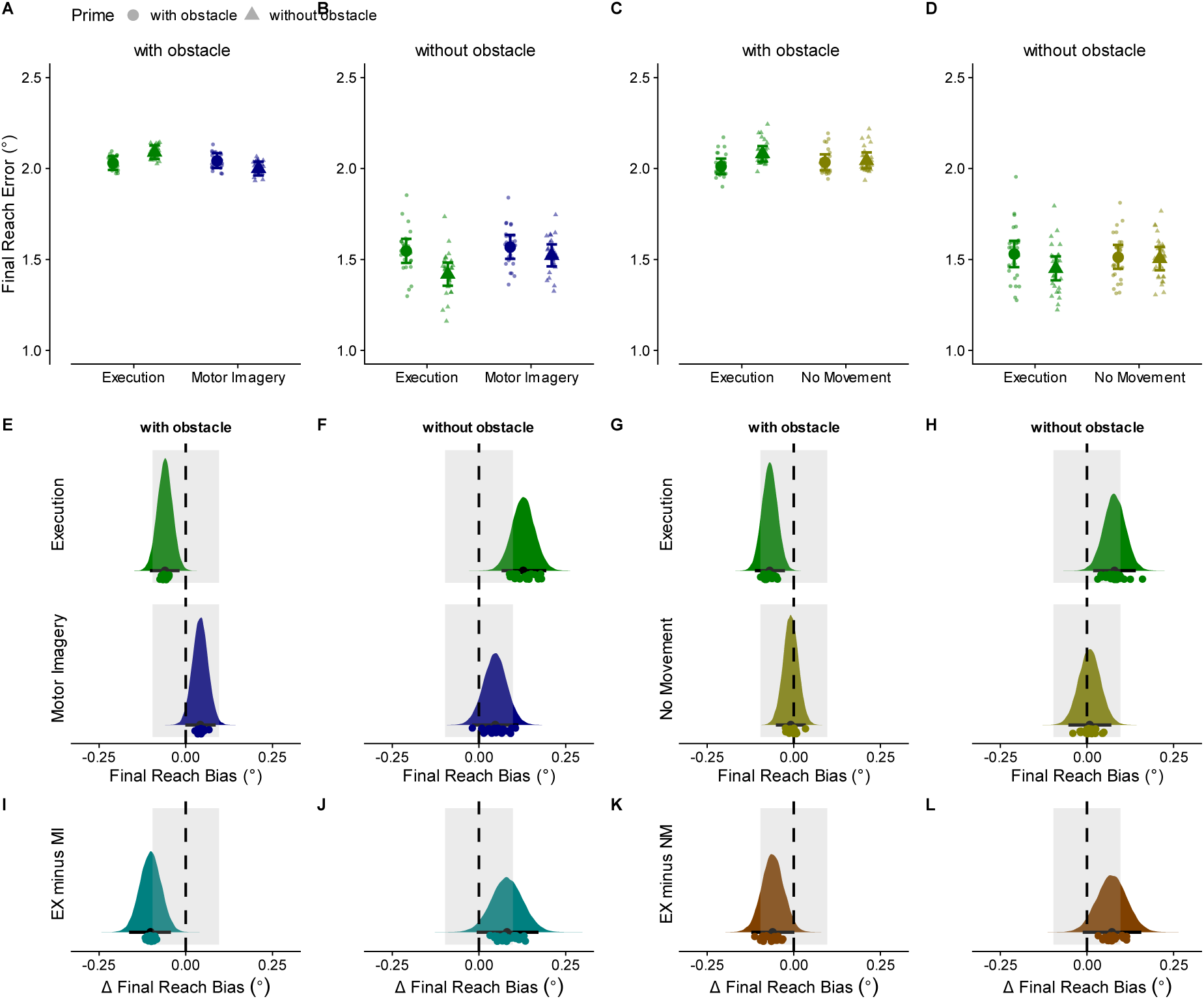
Influences of prime movement on feedback-related movement aspects in Experiment 1 (Motor Imagery) and Experiment 2 (No Movement). **A-D** Final reach error for Experiment 1 (AB) and Experiment 2 (CD). **E-H** Differences in final reach error for probes after primes with vs. without an obstacle for Execution and Motor Imagery (EF) and No Movement (GH). **I-L** Difference of final reach error difference between Execution and Motor Imagery **(IJ**) and No Movement trials (**KL**), respectively. All other details are as in Figure 2.

Thus, the final reach error was modulated only after prime movements had been executed but not when they had been imagined.

The result pattern was similar in Experiment 2 (No Movement). Final reach error was smaller when the prime movement was executed and prime and probe required similar movements (probe with obstacle: difference = -0.07°, [-0.11,-0.03], pd > 99%, 81% in ROPE; probe without obstacle: difference = 0.08°, [0.02,0.14], pd > 99%, 64% in ROPE; Fig. 4GH green data). When the prime movement had been withheld (No Movement condition), no history effect emerged for the final reach error (probe with obstacle: difference = -0.01°, [-0.05,0.03], pd = 66%, 100% in ROPE; probe without obstacle: difference = 0.01°, [-0.05,0.07], pd = 60%, 100% in ROPE; Fig. 4GH yellow data). Direct comparison of execution and non-executed trials confirmed a stronger influence of the previously executed movement, though again only in trials with an obstacle (difference in final error difference = -0.06°, [-0.12,0.00], pd > 97%, 79% in ROPE), but not in trials without an obstacle (difference in final error difference = 0.07°, [-0.01,0.16], pd = 95%, 65% in ROPE; Fig. 4KL).

Thus, history effects are not confined to feedforward-related movement aspects but occur also for feedback-related movement aspects. Critically, feedback-related history effects depend on the overt execution of the prior movement and do not emerge during imagery. This distinction suggests that feedback-related history effects reflect a sensory history, much in contrast to feedforward-related history effects, which are evoked also by imagery and, thus, reflect a motor history.

A potential objection to this conclusion is that the observed differences in final reach error may not truly reflect feedback-based mechanisms but instead may have naturally emerged from differences in the movement trajectories themselves. For example, in obstacle-absent probe trials, the larger final reach error following obstacle-present (compared to obstacle-absent) prime movements (Fig. 4B, green data) could be attributed to the larger initial reach bias. This bias may have produced more curved probe trajectories, causing the target to be approached at a steeper angle and thereby increasing final reach error.

Notably, this explanation is incompatible with the error pattern we observed for obstacle-present probe movements: the final reach error was smaller after prime movements with than without an obstacle (Fig. 4A, green data). This pattern is the opposite of what would be expected if final reach error were simply a byproduct of more curved trajectories. At least three aspects of the current results further support this conclusion. First, if final reach errors were determined primarily by initial reach angle, final reach error should correlate with initial reach angle on a trial-by-trial basis.

Contrary to this prediction, the reduction in final reach error when prime and probe required similar movements was observed across a broad range of initial reach angles and not primarily when the initial reach error was large (Fig. S2A–D). Second, we selected trials with comparable probe initial reach error after primes with and without obstacles (see Supporting Information for details). If the final reach error depended solely on the initial reach error, this data trimming procedure should eliminate any differences between the prime conditions with and without an obstacle; contrary to this prediction, the difference in final reach error (i.e., final reach error was smaller when prime and probe required similar movements) was present also in the trimmed dataset (Fig. S3). Third, we assessed reach precision and accuracy using endpoints measured 150 ms after the cursor entered the target and when its resultant velocity fell below 5 cm/s; we reasoned that such late data points should be least influenced by any potential feedforward influences and amply allow for execution of feedback-induced corrections. Still, the result pattern was similar and showed increased performance (better precision and accuracy following executed prime movements than imagined or no movements; Fig. S4 and S5). In sum, final reach error is modulated independently of the initial reach error, lending support to our proposal that it reflects genuine aspects of movement-related history. This conclusion directly links to the finding that only the initial but not the final reach error is affected when movements are imagined, again emphasizing that the two types of error reflect distinct processes: feedforward-related history effects arise from motor-related processing, whereas feedback-related history effects arise from sensory-related processing.

### Reach bias is similar across target locations and stable across the experiment

Our results contrast with a recent study (Roberts et al., 2025) that employed a similar task but did not observe any bias in initial reach error following motor imagery, leading the authors to suggest that motor imagery does not influence subsequent reach trajectories. This study differed from ours in several respects: Motor imagery and execution trials were administered in separate blocks, the task comprised only one target location, and the number of trials was considerably lower (40 compared to 640). To address whether these procedural differences may be responsible for the diverging result patterns, we ran Experiment 3 as a repeat of Experiment 1 but with prime execution and imagery acquired in separate blocks.

The overall result pattern of Experiment 3 was strikingly similar to that of Experiment 1. A reach bias was evident after both Execution (probe with obstacle: reach bias = 7.9°, [5.5,10.3], pd > 99%, 0% in ROPE; probe without obstacle: reach bias = 15.8°, [12.2,19.6], pd > 99%, 0% in ROPE) and Motor Imagery (probe with obstacle: reach bias = 2.9°, [0.8,5.0], pd > 99%, 35% in ROPE; probe without obstacle: reach bias = 6.7°, [3.8,9.7], pd > 99%, 0% in ROPE), and again larger after execution (probe with obstacle: difference in reach bias = 4.9°, [2.4,7.5], pd > 99%, 0% in ROPE; probe without obstacle: difference in reach bias = 9.1°, [5.5,12.6], pd > 99%, 0% in ROPE; Fig. S6). The bias was present for all target locations (Fig. S7, Table S2), and comparable across the different blocks (Fig. S8, Table S3, see Supporting Information for a detailed description of the results of Experiment 3). Thus, isolating imagery from execution did not affect the emergence of an imagery-based history effect in our experiment.

### Reach bias is temporally short-lived

Since mimicking the experimental approach of Roberts et al. (2025) did not resolve the divergence of their previous and our present results, we next re-analyzed their data in our Bayesian framework, suspecting that our statistical approach may be sensitive to differences overlooked in the original report. Our analysis did not provide any evidence of a history effect after imagined prime movements, confirming the original statistical report (Motor Imagery probe with obstacle: reach bias = -1.6°, [-5.1,1.9], pd = 82%, 66% in ROPE; probe without obstacle: reach bias = 1.8°, [-1.7,5.3], pd = 84%, 63% in ROPE). However, a history effect was also absent after overtly executed primes (probe with obstacle: reach bias = 3.2°, [-0.3,6.7], pd = 96%, 43% in ROPE; probe without obstacle: reach bias = 1.4°, [-2.2,5.1], pd = 78%, 68% in ROPE; Fig. S14). In other words, the design of Roberts et al. did not produce the key marker of the well-known history effect; in light of a failure to replicate this effect, the absence of an imagery effect is difficult to interpret.

We were surprised by Roberts et al.’s failure to obtain a history effect, not only given the rather extensive previous literature that has documented the existence of the phenomenon, but also because in our present study, the effect was present in every single participant. Therefore, we scrutinized their data further. Their median RT was considerably longer than in any of our experiments (Roberts et al., 2025: Execution: 379 ms, Motor Imagery: 520 ms; our Experiment 1: Execution = 295 ms, Motor Imagery = 301 ms; Experiment 2: Execution = 294 ms, No Movement = 339 ms; Experiment 3: Execution = 272 ms, Motor Imagery = 326 ms; Fig. 5). This led us to test whether the reaction time of a given trial affected the emergence of the initial reach bias. Indeed, for both execution and imagery trials in our Experiments 1 and 3, the reach bias was most pronounced in trials with short response latency and declined with increasing RT, indicating that the history effect was temporally short-lived (Fig. 6). In fact, the reach bias was absent (or largely reduced) around RTs of 350-400ms in our experiments, an RT that lies below the average RT of Roberts et al. (2025).

**Figure 5.**
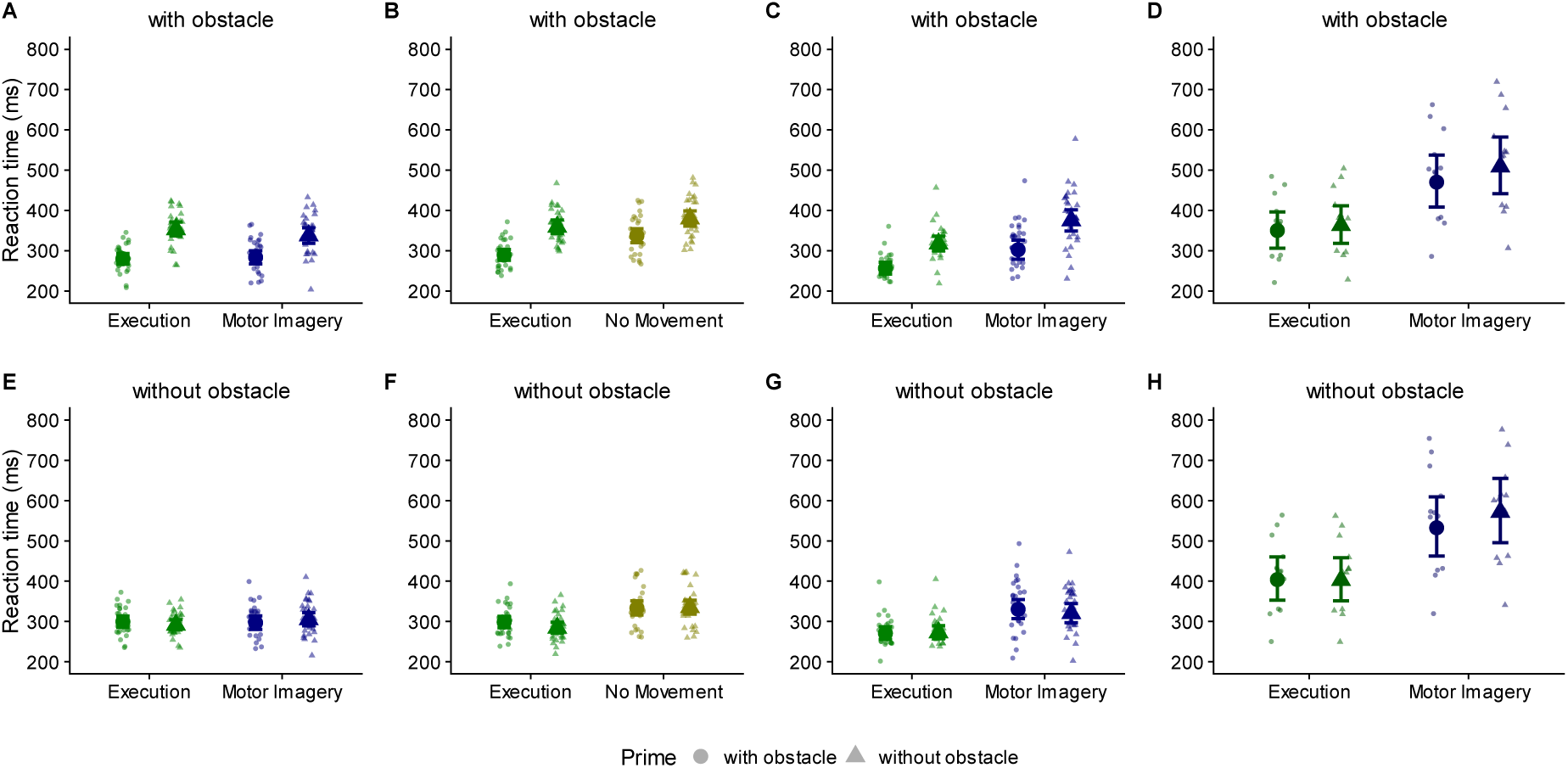
Reaction times (in ms) as a function of prime phase, probe phase, and trial type for Experiment 1 (**AE**), Experiment 2 (**BF**), Experiment 3 (**CG**), and Roberts et al. 2025 (**DH**). Large symbols represent the group estimates and error bars the 95% highest-density interval (HDI). Small symbols represent participant estimates.

**Figure 6.**
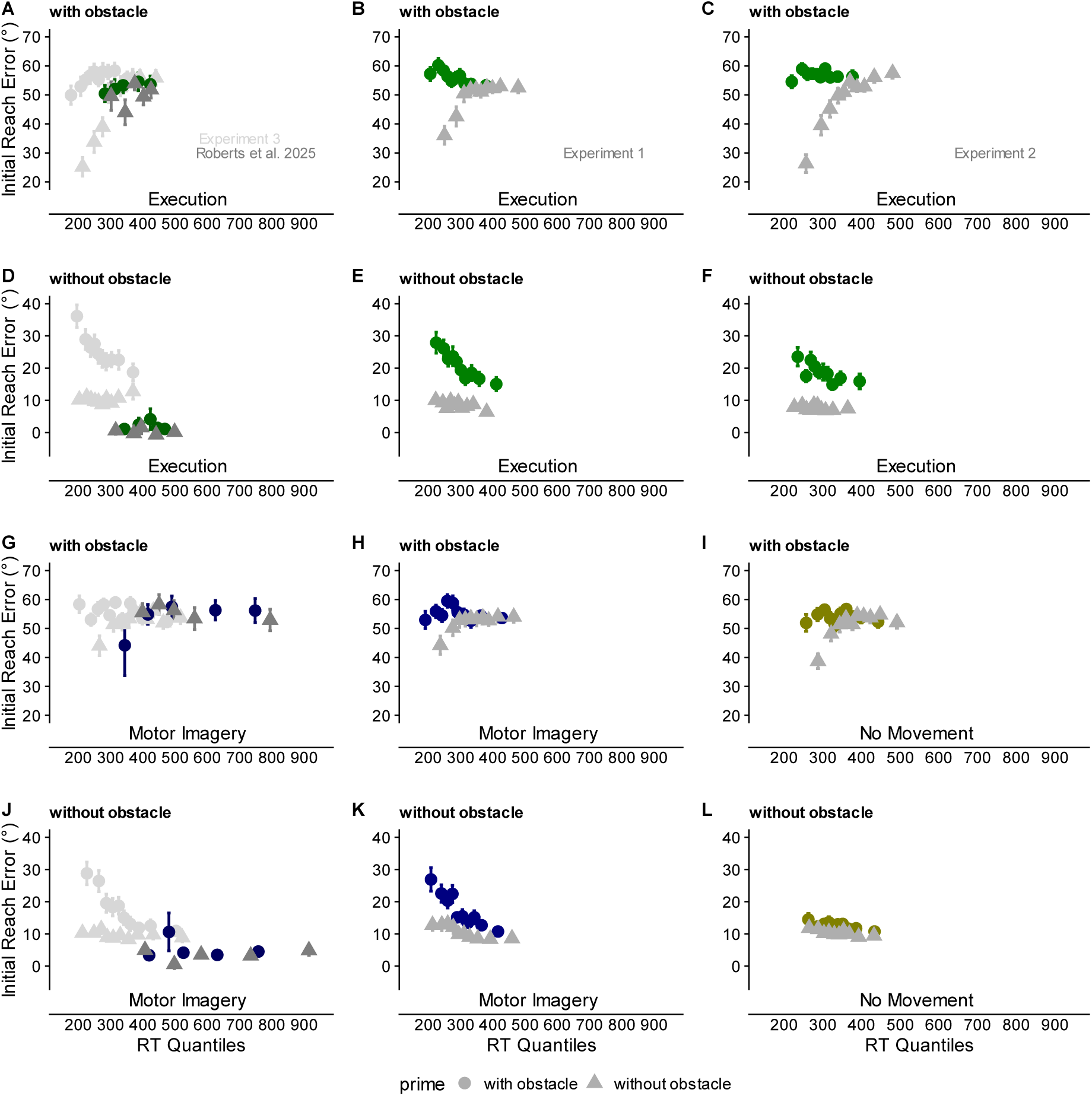
Influences of prior movement on feedforward aspects are temporally short-lived. Initial reach error as a function of reaction time. RTs were vincentized into equal-sized quantiles (deciles for Experiments 1-3; quintiles for data from Roberts et al. 2025) separately for each participant and experimental condition. Group average (±1 s.e.) values for Initial Reach Error corresponding to reach reaction time quantile. **ADGJ** Data from Roberts et al. 2025 (dark dots) and Experiment 3 (light dots). **BEHK** Data from Experiment 1. **CFIL** Data from Experiment 2.

Fitting with this observation in our dataset, no temporal decline was present in their data (Fig. 6). Thus, we contend that the study of Roberts et al. should not be interpreted as evidence against the ability of motor imagery to induce motor history effects, both because it did not establish a history effect for executed trials in the first place, and because their null findings are consistent with our finding that history effects unfold primarily for probe trials that are initiated fast.

## Discussion

Despite the wealth of evidence demonstrating that motor performance can be biased by prior movements (Diedrichsen et al., 2010; Schütz et al., 2011; Seegelke et al., 2015, 2013; Tsay et al., 2022; Verstynen and Sabes, 2011), it has remained open whether such biases are primarily driven by sensory or motor aspects of the movement. We tested whether history effects obtained following overt, physically executed movements can also be elicited by imagined movements, with the latter involving the motor but not the sensory aspects of overt movements. Furthermore, we extended the common approach of investigating history effects in early, feedforward, planning-related to also examine late, feedback-related movement aspects. Early, feedforward aspects were influenced by a history established both by overt and imagined movement, suggesting a motor rather than sensory origin. In contrast, late, feedback-related aspects were modified only by a history established by overt, physically executed movements, suggesting that history effects for the later part of the movement reflect sensory rather than motor factors of previous movements. Accordingly, motor and sensory signals make distinct contributions to motor history and shape specific aspects of current motor performance.

An important aspect of our findings is that imagery-based history effects were smaller than those obtained after overt execution. Speculatively, this difference may be attributable to the need for an inhibitory mechanism that suppresses movement during imagery. During both motor imagery and regular motor planning, cortical processing must not activate descending efferent pathways to prevent overt execution. Accordingly, the two cases may rely on a common inhibitory mechanism.

Research on inhibition has often employed stop-signal tasks, which measure how quickly and effectively an already-prepared action can be canceled when it must be stopped unexpectedly (Logan and Cowan, 1984; Verbruggen and Logan, 2008). We observed a reduction of history effects similar to that reported in the present study when participants were required to cancel a planned prime movement in response to a stop-signal cue (Seegelke and Heed, 2026). In contrast, when the hand was blocked and prime movement execution was physically prevented – without participants having to actively inhibit the movement – history effects were comparable to those following normally executed primes. Thus, history effects are reduced when preventing prime execution involves active inhibition, but they remain unchanged when execution is prevented by external means such as physical blocking. Thus, inhibitory control appears to act on the neural activity responsible for motor planning and motor imagery (Rieger et al., 2024), and it is this interference we propose attenuates the functional impact that planning- and imagery-related activity would otherwise have on the build- up of a motor history.

When we removed all motor-related activity in the prime phase, a small history effect remained: even though participants in our No Movement experiment could just wait for the instructional cue for the probe movement, simply seeing the target and obstacle resulted in a small but credible reach bias for the probe movement. It is conceivable that the residual bias reflects visuomotor priming. It is well established that the mere visual presentation of an object can decrease response latencies of subsequent actions afforded by the object (Craighero et al., 1996; Masson et al., 2011) and bias movement kinematics (Hesse et al., 2008; Seegelke et al., 2016). Likewise, reaches to a target location can be biased towards concurrently presented distractor targets (Moher and Song, 2013; Tipper et al., 1992). Thus, the visual context during the prime phase, such as the target configuration and the presence of an obstacle, apparently primes appropriate actions. This interpretation is reinforced by the reduced probe response latencies when an obstacle was present for both prime and probe (observed for Experiments 1-3, see Fig. 6). This small effect notwithstanding, the largest part of the history effect we observed was attributable to motor-related processing. The overall pattern of visual-sensory aspects playing a role in movement history while not being its key driver is consistent with previous evidence: whether vision of the target layout and hand cursor was provided or prohibited left history effects largely unaffected (Glover and Dixon, 2013; Seegelke and Heed, 2026). Thus, not only is the influence of vision comparatively small, it is also not required for history effects to occur.

Previous studies have not usually investigated potential history effects in those phases of the movement in which sensory feedback can exert an influence. History effects were present even at the very end of the movement, that is, for aspects related to target acquisition. Critically, the comparison of execution and imagery as determinants of movement history revealed a dissociation between feedforward and feedback contributions to movement history: overtly executed movements evoked history effects both in feedforward and feedback-related movement aspects, whereas motor imagery affected only feedforward but not feedback-related movement aspects.

We note that the effects on feedback-based movement aspects were small and, overall, less statistically reliable than feedforward-related effects. For instance, a credible modulation of final reach error by overtly executed primes was present in all three experiments, but endpoint precision and accuracy were credibly affected only in Experiment 1. Critically, performance was consistently superior after overtly executed than after imagined primes, both regarding precision and accuracy. In our setup, the small target (1 cm in diameter) imposed high endpoint precision demands, which may have prevented stronger manifestation of feedback-related aspects. On the other hand, the small target size forced all trajectories to end in a highly confined spatial area, which we reasoned would maximize final reach error, as it forces all trajectories to curve into this very same spot. Given that previous research on motor history has not focused on target-related movement aspects, there is likely room to optimize its experimental assessment.

Prevalent motor control theories argue that feedforward and feedback processes are governed by a common neural circuitry (Pruszynski, 2014; Scott, 2016). For instance, in sensorimotor adaptation paradigms, learning of new feedforward commands transfers to feedback responses (Cluff and Scott, 2013; Maeda et al., 2018; Wagner and Smith, 2008) and vice versa (Maeda et al., 2020). Yet not all evidence is consistent with this view. For instance, in a mirror-reversed reaching task, feedforward control adapted even when online correction during reaches, and thus, obtaining the respective sensory information, was prevented. In contrast, feedback control adapted only when online sensory information was available (Kasuga et al., 2015). Both the relevance of sensory information and the separate tuning for feedforward and feedback are consistent with our current findings. In this scheme, motor imagery engages predictive internal forward models (Kilteni et al., 2018) and biases movement planning and initial movement estimates, thus affecting feedforward control. However, because motor imagery lacks sensory feedback, it does not recruit those feedback control mechanisms that process ongoing sensory input to adjust movement online.

In conclusion, our experimental findings suggest that distinct processing exists for motor history of preparatory versus execution-related activity: planning-related history is sufficient to bias initial reach direction, whereas execution-related sensory consequences are required to adjust feedback-based control. Consequently, motor imagery leaves a “planning trace” that influences the next movement’s feedforward component, but only actual execution leaves the “feedback trace” that improves endpoint accuracy and precision. This division underscores that the sensorimotor system leverages different information channels—efferent plans versus afferent consequences—to sculpt distinct facets of subsequent performance.

## Methods

### Participants

We recruited a total of 93 participants (Experiment 1: N = 32, Experiment 2: N = 31, Experiment 3: N = 30). For Experiment 1, we excluded data from one participant who did not complete the experiment. For Experiment 2, we excluded data from one participant who did not follow the instructions. Thus, Experiment 1 comprised 31 individuals (15 female, 16 male, mean age = 23.0 years, SD = 3.48, range = 18-35), Experiment 2 comprised 30 individuals (18 female, 12 male, mean age = 24.4 years, SD = 6.06, range = 18-50), and Experiment 3 comprised 30 individuals (27 female, 3 male, mean age = 22.8 years, SD = 1.77, range = 20-26). Sample size was comparable to our similar study (Seegelke and Heed, 2026) and substantially higher compared to previous studies (Jax and Rosenbaum, 2007; Roberts et al., 2025) and was determined based on our resources in terms of acquisition time. All participants had normal or corrected-to-normal vision and received course credit for their participation. The University of Salzburg Ethics Committee approved all procedures (Ethical Application Ref: EK-GZ 32/2023).

### Apparatus and tasks

Participants operated an optical computer mouse (MS116, Dell Inc.) with their dominant hand to perform center-out movements with the cursor from a central start location to one of 8 target locations. The targets were positioned on an imaginary circle at 22.5°, 67.5°, 112.5°, 157.5°, 202.5°, 247.5°, 292.5°, and 337.5°, 8 cm from the start location. In a subset of trials, an obstacle appeared between start and target location which had to be circumnavigated. Stimuli were presented on a 25” computer monitor (AW2523HF, Dell Inc.; 1920x1080 pixels; 240Hz refresh rate) positioned at a viewing distance of about 50 cm. Start and target locations were white circles (1 cm in diameter). Obstacles were white ellipses (major axis = 2 cm, minor axis = 1 cm) and were positioned halfway between start and target location with the major axis oriented perpendicular to a line connecting start and target location. Stimulus presentation and response recording were controlled with PsychoPy v2023.2.3 (Peirce et al., 2019).

### Procedure

Participants performed two movements in succession, a prime followed by a probe reaching movement (Fig. 1A). Each trial started with the presentation of the start location at the center of the screen. Participants then moved the cursor into this location and maintained it there for 500 ms. During execution trials, the prime target then appeared at one of the 8 target locations, and participants moved the cursor to the respective target location. After the cursor was maintained at the target location for 250 ms, the target location disappeared, and the participant moved the cursor back to the start location. After maintaining the cursor there for 500 ms, the probe target appeared (at the same location as in the prime), and participants moved the cursor to the target location. After the cursor was maintained at the location for 250 ms, the participant then moved the cursor back to the start location triggering initiation of the next trial. During Motor Imagery trials (Experiments 1 and 3), the target location was displayed in red (imagery-signal) for 500 ms, and participants were instructed to withhold any overt movement and to imagine the cursor movement from start to target and back. They were asked to signal when their imagined movement reached the start location again by pressing the left mouse button. After participants pressed the mouse button, the probe phase was initiated which was identical to the probe phase of execution-trials. During No Movement trials (Experiment 2), the target location was displayed in red (as in Motor Imagery trials) for 500 ms, signaling participants to withhold any overt movement. In contrast to the Motor Imagery trials however, they were not instructed to imagine the movement. After participants maintained the cursor at the start location for 2000 ms (to approximately match the duration between prime target onset and probe target onset of execution trials), the probe phase was initiated which was identical to the probe phase of execution trials. In a subset of trials, an obstacle could appear either during the prime phase, the probe phase, or both. In obstacle-present trials, the obstacle appeared simultaneously with the target location and participants were instructed to move (or to imagine moving) the cursor around the obstacle to reach the target location. The obstacle remained present also during the back-to-start movements and was extinguished once participants moved the cursor back in the start location.

If participants did not move the cursor out of the start position within 750 ms after target onset, the message “Respond faster” was shown on the screen in black letters for 1000 ms. Participants still had to reach to the target location and back to the start location to initiate the next phase/ trial. If participants hit the obstacle, it turned red for 500 ms and the message “You shall not pass” was displayed on the screen in black for 1000 ms. If participants moved the cursor out of the start location during a Motor Imagery/No Movement trial, the message “Stop-Signal: Don’t move” was displayed on the screen in black letters for 1000 ms, and there was a delay of 1500 ms before participants could initiate the probe phase.

In each trial, we manipulated whether an obstacle was present or absent during the prime and/or probe phase. Half of the trials were execution trials, the other half imagery trials, and movements had to be directed to each of the 8 target locations; target location was always identical for prime and probe movement. Thus, there were 64 conditions in total, comprised of the factors obstacle prime (present, absent) x obstacle probe (present, absent) x trial type (execution-trial, imagery-trial) x target location, i.e. a 2 x 2 x 2 x 8 design.

In Experiments 1 and 2, Execution and Motor Imagery/No Movement trials were presented within the same block. Participants performed 11 blocks, each consisting of 64 trials. In each block, each condition was presented once, and the order of presentation was randomized. The first block was considered practice and not analyzed. In Experiment 3, Execution and Motor Imagery trials were presented in separate blocks. Participants performed 11 execution blocks followed by 11 imagery blocks (or vice versa, counterbalanced across participants), each consisting of 32 trials. Each remaining condition was presented once, and the order of presentation was randomized. The first block was considered practice and not analyzed. Prior to Experiments 1 and 3, participants completed the MIQ-RS motor imagery questionnaire (Gregg et al., 2010), which assesses individuals’ visual and kinesthetic imagery abilities using seven different and relatively simple movements on a 7-point Likert scale ranging from 1 (= very hard to see/feel) to 7 (= very easy to see/feel). Further, participants evaluated their motor imagery after each block, rating the ease with which they imagined the movements on a 7-point Likert scale ranging from 1 (very hard to imagine) to 7 (= very easy to imagine), and how frequently they imagined the movements on a 3-point scale (1 = in fewer than half of the trials, 2 = in most trials, 3 = in every trial; Sheahan et al., 2018). Each experiment took about 1.5 hours.

### Data processing and analysis

We processed kinematic data with custom-written scripts in MATLAB (version R2023a; The MathWorks, Natick, MA). We filtered kinematic data using a second-order butterworth filter with a cutoff frequency of 10 Hz. We determined reach onset as the time of the sample in which the vectorial velocity exceeded 50 mm/s and reach offset as the time of the sample in which the cursor entered the target location of the respective phase (i.e., prime and probe). We defined reaction time (RT) as the time between target onset and reach onset, and movement time (MT) as the time between reach onset and offset. We defined initial reach error as the absolute angular deviation between the vector from the cursor position at reach onset to the cursor position when it first left the start location and the vector from the cursor position at reach onset to the target center. The average time between movement onset and the timepoint when the cursor left the start location was considerably below the time necessary to utilize visual feedback (∼100 ms; Scott, 2016); Experiment 1: prime movement mean = 36 ms, SD = 10, probe movement mean = 43 ms, SD = 12; Experiment 2: prime movement mean = 36 ms, SD = 10, probe movement mean = 43 ms, SD = 11; Experiment 3: prime movement mean = 37 ms, SD = 12, probe movement mean = 44 ms, SD = 14). Hence, the extracted initial reach direction reflects a readout of a pure feedforward motor command.

We used three measures to consider feedback-related effects: First, we quantified final reach error as the absolute value of the angular deviation between the vector connecting cursor position at reach onset and the target center and the vector connecting cursor position at reach onset and offset/target hit. While this measure might not provide a readout of pure feedback-related aspects (see Results section), it is extracted at a timepoint where it certainly is subject to corrective processes; median MT was 388 ms, 388 ms, and 434 ms for Experiments 1, 2, and 3, respectively. Second, as a measure of reach precision, we obtained reach endpoints as the cursor position 150 ms after it entered the target and when its resultant velocity was below 5 cm/s and calculated the area of the 95% confidence ellipse (in mm²) for these reach endpoints. Third, as a measure of reach accuracy, we calculated absolute reach endpoint error as the Euclidean distance (in mm) of the reach endpoints to the target center. We chose the combined duration/velocity criterion to minimize any potential conflating influence of feedforward processes and/or speed-accuracy tradeoffs on terminal feedback. For Motor Imagery trials, the duration of the imagined movements (during the prime phase) was calculated as the time from prime target onset to button press. As an equivalent in Execution trials, we calculated the duration of prime movements as the sum of RT, MT, and the movement duration for the movement back from target to start position. We excluded trials in which participants moved the cursor out of the start location during a Motor Imagery/No Movement trial (Experiment 1: N = 773, 7.8% of imagery trials; Experiment 2: N = 358, 3.7% of no movement trials; Experiment 3: N = 374, 3.9% of imagery trials), trials in which they reacted too slowly during the prime (Experiment 1: N = 152; Experiment 2: N = 47; Experiment 3: N = 35) or probe phase (Experiment 1: N = 168; Experiment 2: N = 81; Experiment 3: N = 289), trials in which they hit the obstacle during the prime (Experiment 1: N = 161; Experiment 2: N = 89; Experiment 3: N = 176) or probe phase (Experiment 1: N = 296; Experiment 2: N = 264; Experiment 3: N = 280), trials in which movement onset could not be detected during the prime (Experiment 1: N = 127; Experiment 2: N = 144; Experiment 3: N = 426) or probe phase (Experiment 1: N = 198; Experiment 2: N = 178; Experiment 3: N = 482), and trials which did not exhibit smooth trajectories (i.e., movements with more than two velocity peaks; Experiment 1: N = 806; Experiment 2: N = 666; Experiment 3: N = 771). This led to exclusion of 2,479 trials (12.5%) in Experiment 1, 1,756 trials (9.1%) in Experiment 2, and 2,632 in Experiment 3 (13.7%). Finally, we removed trials in which RT was shorter than 100 ms or MT longer than 1000 ms (Experiment 1: N = 471; Experiment 2: N = 268; Experiment 3: N = 717).

### Statistical approach

We fit Bayesian regression models created in Stan (http://mc-stan.org/) and accessed with the package brms version 2.22 (Bürkner, 2017) in R (R Core Team, 2025). We fit separate models for each of our dependent variables: Initial reach error of probe movement, final reach error of probe movement, reach end point variability of probe movements, reach end point error of probe movement, reaction time (RT) of probe movement, and prime duration. For RT, we included the within-subject categorical variables prime phase (obstacle, no obstacle), probe phase (obstacle, no obstacle), trial type (Execution, Motor Imagery/No Movement), and all interactions as independent predictors. For prime duration, we included the within-subject categorical variables prime phase (obstacle, no obstacle) and trial type (Execution, Motor Imagery/No Movement), and all interactions as independent predictors. We modeled RT and prime duration using shifted log-normal distributions.

Distributions of initial reach error and final reach error differed considerably between trials with or without an obstacle during the probe phase. We thus fit separate models for each probe condition (with obstacle, without obstacle) with the within-subject categorical variables prime condition (with obstacle, without obstacle), trial type (Execution, Motor Imagery/ No Movement), and all interactions as independent predictors. We modeled Initial reach error using truncated gaussian and log-normal distributions (lower bound [lb]: 0°; upper bound [ub]180°) for trials with and without an obstacle in the probe phase, respectively. We modeled Final reach error using skewed gaussian and truncated (lb: 0°, ub: 4°) gaussian distributions for trials with and without an obstacle in the probe phase, respectively. For end point variability (i.e., precision) and end point error (i.e., accuracy), we included the within-subject categorical variables prime phase (obstacle, no obstacle), probe phase (obstacle, no obstacle), trial type (Execution, Motor Imagery/ No Movement), and all interactions as independent predictors and we used truncated (precision: lb: 0mm²; accuracy: lb: 0mm, ub: 5mm) gaussian distributions. We set orthogonal contrasts using the set_sum_contrast() command in afex version 1.3- 0 (Singmann et al., 2020). For all models, we included random intercepts and slopes for all main effects and interactions, and specified weakly informative priors to regularize parameter estimates and improve convergence. For re-analysis of the data of Roberts et al. (2025), we included the within-subject categorical variables prime phase (obstacle, no obstacle), probe phase (obstacle, no obstacle), trial type (Execution, Motor Imagery), and all interactions as independent predictors. For initial reach error, we used a gaussian distribution, for RT we used a shifted log-normal distribution. Due to the limited number of trials, we restricted the random-effects structure to random intercepts to ensure stable estimation and model convergence.

For all analyses, the estimation of parameters’ posterior distributions was obtained by Hamiltonian Monte-Carlo sampling with 4 chains, 1,000 sample warmup, and 11,000 iterations and checked visually for convergence (high ESS and Rhat ≈ 1). Exact model specifications for each model can be obtained from the code available in the corresponding author’s github repository (https://github.com/CSeegelke1/Motor-History-in-Motor-Imagery). We used the package bayestestR version 0.13.1 (Makowski et al., 2019a, 2019b) to describe the parameters of our models. We report the median as a point estimate of centrality and the 95% credible interval (CI) computed based on the highest-density interval (HDI) to characterize the uncertainty related to the estimation. Following current recommendations for Bayesian inference (Kruschke, 2018; Makowski et al., 2019a, 2019b), we distinguish between the *existence* and *practical significance* of an effect, as these address conceptually different questions and need not yield the same conclusion. Effect existence — that is, whether an effect is credibly different from zero in a particular direction—was assessed using the probability of direction (*pd*), defined as the proportion of the posterior distribution sharing the sign of the posterior median. Following the reference values proposed by Makowski et al. (2019b), we interpreted *pd* ≤ 95% as *uncertain*, *pd* > 95% as *possibly existing*, *pd* > 97% as *likely existing*, and *pd* > 99% as *probably existing*. Practical significance—that is, whether an effect was sufficiently large to be considered non-negligible—was assessed using a region of practical equivalence (ROPE; Kruschke, 2018) of ±0.1 effect sizes around zero and by calculating the proportion of the highest-density interval (HDI) falling within this region. Effects whose HDI fell entirely outside the ROPE were considered practically significant (i.e., non-negligible), whereas effects whose HDI fell entirely within the ROPE were considered practically negligible (i.e., practically equivalent to zero). Partial overlap between the HDI and ROPE was interpreted as inconclusive with respect to practical significance. Because *pd* and ROPE-based inference address different questions, an effect may show strong evidence for existing in a particular direction while remaining too small to be practically meaningful. We therefore report and interpret both indices jointly for every effect rather than treating either measure as a binary indication of whether an effect is simply “present” or “absent.”

## Data and code availability

The preprocessed data required to reproduce the analyses are available on Zenodo (https://doi.org/10.5281/zenodo.22251643). Analysis scripts are available on GitHub (https://github.com/CSeegelke1/Motor-History-in-Motor-Imagery) and archived on Zenodo (https://doi.org/10.5281/zenodo.22252677). The data and software records are cross-linked through their metadata.

