## Supplementary Materials for "Distinct contributions of motor imagery and execution to history-dependent biases in reaching"

**Table S1.** Correlations between Reach Bias (i.e., difference in initial reach error between trials with an obstacle in the prime movement and trials without an obstacle in the prime movement) and Motor Imagery Questionnaire Scores in Experiment 1.

| Measure | Trial Type | Probe | r | HDI | pd (%) |
|---|---|---|---|---|---|
| MIQ Score Overall | Execution | without obstacle | 0.06 | [-0.26, 0.36] | 64.10 |
| MIQ Score Overall | Motor Imagery | without obstacle | 0.16 | [-0.16, 0.46] | 81.30 |
| MIQ Score Overall | Execution | with obstacle | 0.21 | [-0.11, 0.49] | 89.60 |
| MIQ Score Overall | Motor Imagery | with obstacle | 0.23 | [-0.11, 0.51] | 92.55 |
| MIQ Score Visual | Execution | without obstacle | 0.01 | [-0.32, 0.31] | 51.88 |
| MIQ Score Visual | Motor Imagery | without obstacle | 0.07 | [-0.28, 0.38] | 66.17 |
| MIQ Score Visual | Execution | with obstacle | 0.10 | [-0.22, 0.43] | 72.88 |
| MIQ Score Visual | Motor Imagery | with obstacle | 0.29 | [-0.02, 0.56] | 96.62 |
| MIQ Score Kinesthetic | Execution | without obstacle | 0.09 | [-0.22, 0.40] | 69.58 |
| MIQ Score Kinesthetic | Motor Imagery | without obstacle | 0.18 | [-0.15, 0.48] | 85.50 |
| MIQ Score Kinesthetic | Execution | with obstacle | 0.25 | [-0.08, 0.52] | 92.62 |
| MIQ Score Kinesthetic | Motor Imagery | with obstacle | 0.15 | [-0.19, 0.45] | 80.67 |
| Ease of Imagery | Execution | without obstacle | 0.10 | [-0.21, 0.41] | 71.65 |
| Ease of Imagery | Motor Imagery | without obstacle | 0.14 | [-0.19, 0.44] | 81.00 |
| Ease of Imagery | Execution | with obstacle | 0.11 | [-0.20, 0.40] | 73.17 |
| Ease of Imagery | Motor Imagery | with obstacle | -0.07 | [-0.37, 0.27] | 66.67 |
| Count of Imagery | Execution | without obstacle | -0.12 | [-0.42, 0.19] | 75.75 |
| Count of Imagery | Motor Imagery | without obstacle | 0.03 | [-0.31, 0.32] | 57.48 |
| Count of Imagery | Execution | with obstacle | -0.03 | [-0.35, 0.27] | 56.12 |
| Count of Imagery | Motor Imagery | with obstacle | -0.23 | [-0.53, 0.08] | 91.80 |

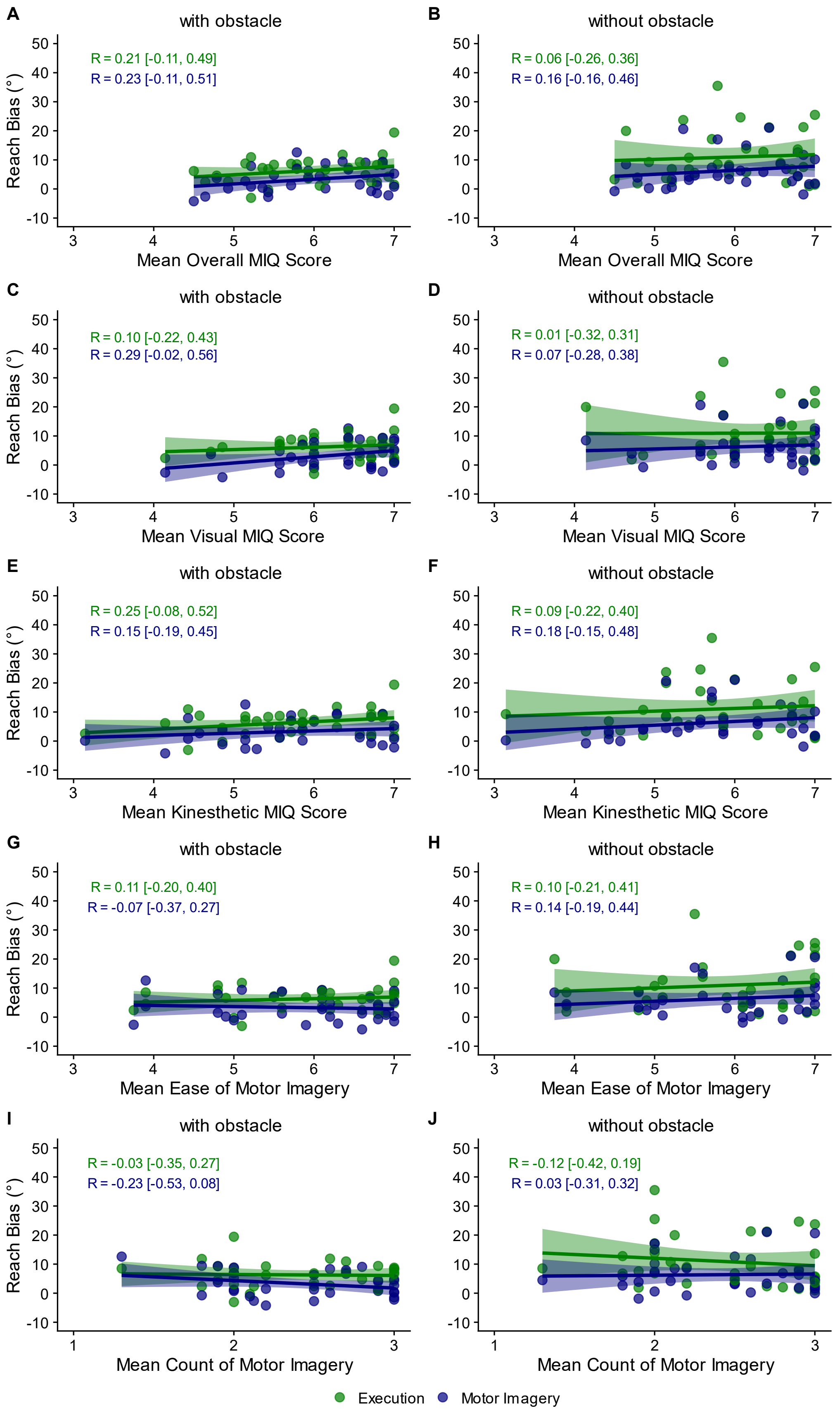


**Figure S1.** Experiment 1. Correlations between Reach Bias and **AB** Mean Overall MIQ Score, **CD** Mean Visual MIQ Score, **EF** Mean Kinesthetic MIQ Score, **GH** Mean Ease of Motor Imagery Score, and **IJ** Mean Count of Motor Imagery Score. Solid lines: linear fit; shading: ±1 s.e. of the fit.

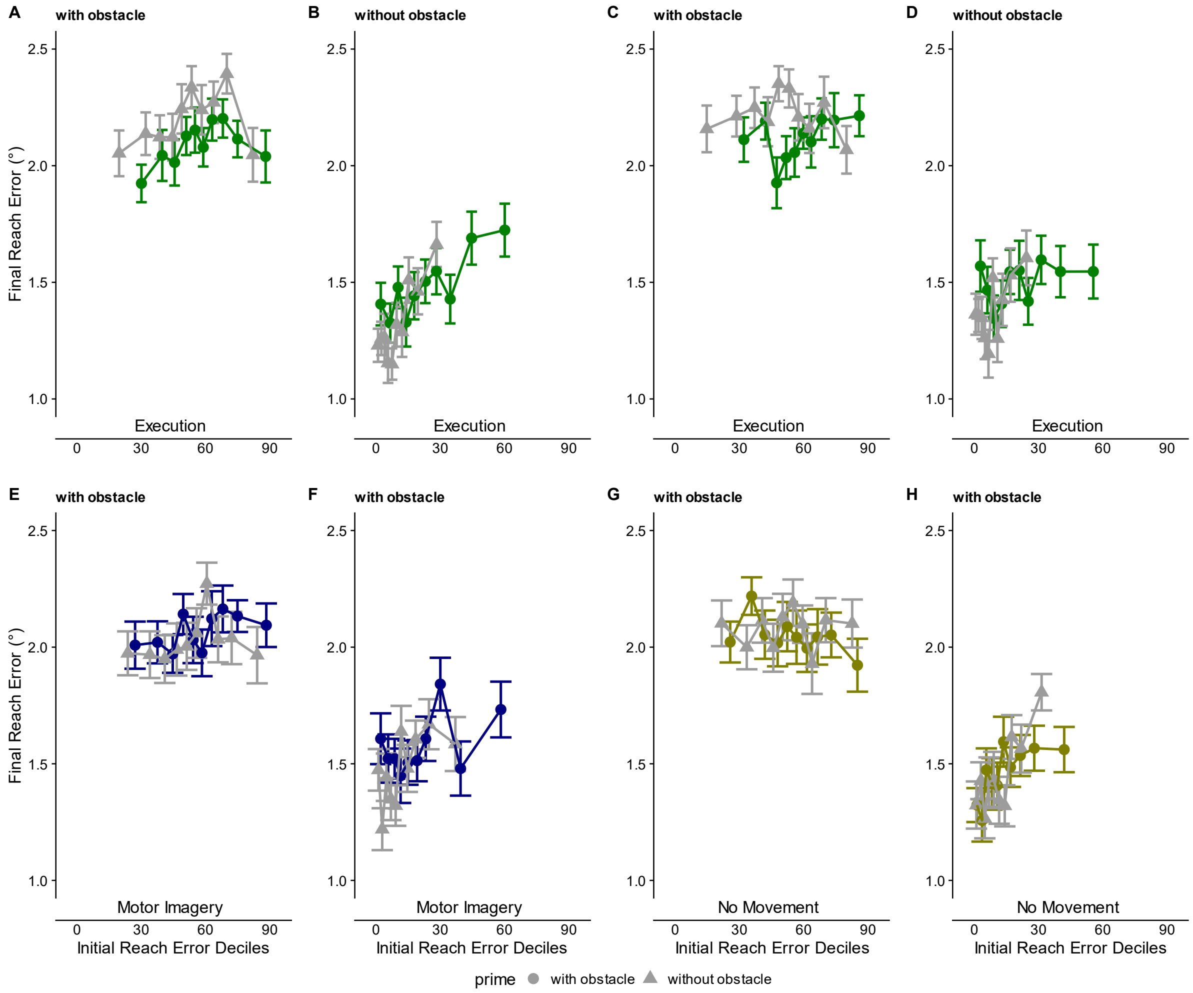


**Figure S2.** Final Reach Error as a function of Initial Reach Error. **ABEF** Experiment 1 (Motor Imagery). **CDGH** Experiment 2 (No Movement). Initial Reach Errors were vincentized into equal-sized deciles separately for each participant and experimental condition. Group average (±1 s.e.) values for Final Reach Error corresponding to Initial Reach Error decile.

### Trimming procedure to mitigate differences in Initial Reach Error

To assess whether the observed differences in final reach error truly reflect feedback-based mechanisms and do not result from differences in the movement trajectories themselves, we applied a trimming procedure separately for Experiment 1 and Experiment 2. To reduce differences in Initial Reach Error between the prime conditions with and without an obstacle, for each participant × probe (with obstacle, without obstacle) × trial type (Execution, Motor Imagery/No Movement), extreme trials were iteratively removed to balance condition means. Specifically, the mean of Initial Reach Error was calculated for each prime level (with obstacle, without obstacle) within each participant × probe × trial type combination. The trial farthest from the overall target mean was removed from the condition with the larger deviation, repeating until the absolute difference between condition means was ≤ 0.05° or a minimum of 40 trials per condition was reached. This procedure ensured within-subject balance across conditions while preserving an adequate number of trials. 4,022 trials (23.8% of the data in Experiment 1) and 4,077 trials (23.7% of the data in Experiment 2) were removed in the trimming process. As intended, data trimming led to a substantial reduction in Initial Reach Bias (Experiment 1: Pre minus Post = 4.3° and 10.7° for Execution trials with and without an obstacle in the probe phase, respectively; Pre minus Post = 1.7° and 5.3° for Motor Imagery trials with and without an obstacle in the probe phase, respectively; Experiment 2: Pre minus Post = 6.7° and 10.5°

for Execution trials with and without an obstacle in the probe phase, respectively; Pre minus Post = 1.6° and 2.0° for No Movement trials with and without an obstacle in the probe phase, respectively), leaving only small differences in Initial Reach Error between trials with and without an obstacle in the prime phase (Experiment 1: Execution probe with obstacle: 2.3°, [1.3,3.3]; Execution probe without obstacle: 2.2°, [0.7,3.7]; Motor Imagery probe with obstacle: 1.4°, [0.4,2.4]; Motor Imagery probe without obstacle: 1.8°, [0.3,3.3]; Experiment 2: Execution probe with obstacle: 2.1°, [1.2,3.1]; Execution probe without obstacle: 2.1°, [0.5,3.8]; No Movement probe with obstacle: 0.6°, [-0.3,1.6]; No Movement probe without obstacle: 0.4°, [-0.5,1.4]). Critically, despite this considerable change in Initial Reach Error, this procedure did not eliminate the differences in Final Reach Error (Fig. S3).

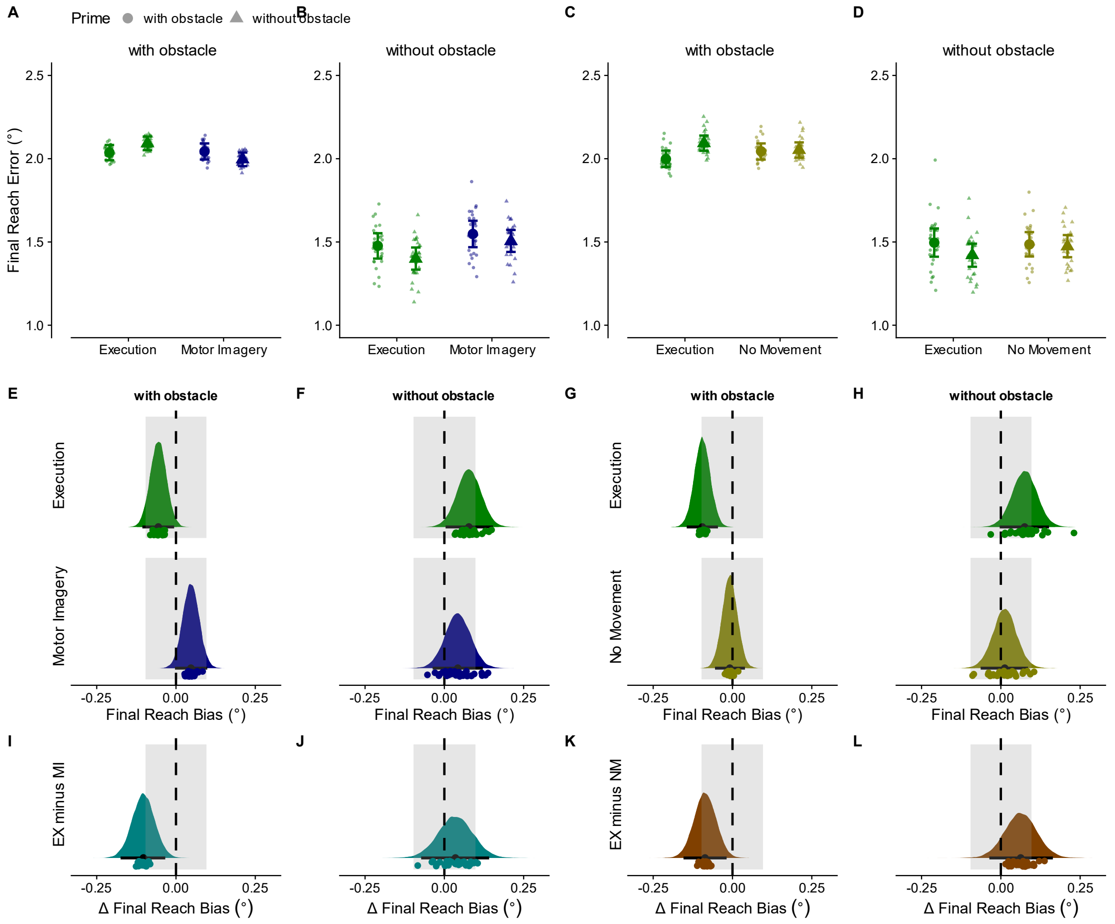


**Figure S3.** Influences of prior movement on feedback aspects in Experiment 1 (Motor Imagery) and Experiment 2 (No Movement) based on trimmed data. Extreme trials were iteratively removed within each participant so that the mean of Initial Reach Error was balanced across prime conditions. Trimming stopped once the condition means differed by ≤ 0.05° or the minimum of 40 trials per condition was reached. **A-D** Final Reach Error for Experiment 1 (AB) and Experiment 2 (CD). Large symbols represent the group estimates and error bars the 95% highest-density interval (HDI). Small symbols represent participant estimates. **E-H** Differences in Final Reach Error between trials with and without an obstacle in the prime movement for Execution and Motor Imagery/ No Movement conditions with and without an obstacle in the probe phase. In Experiment 1, Final Reach Error was smaller when prime and probe necessitated similar movements (probe with obstacle: difference = -0.06°, [-0.11,-0.01], pd = 99%, 90% in ROPE; probe without obstacle: difference = 0.08°, [0.00,0.15], pd = 98%, 63% in ROPE), whereas there was no statistical difference in Motor Imagery (probe with obstacle: difference = 0.05°, [0.00,0.10], pd < 97%, 98% in ROPE; probe without obstacle: difference = 0.04°, [-0.04,0.12], pd = 86%, 85% in ROPE). Likewise, in Experiment 2, Final Reach Error was smaller when prime and probe necessitated similar movements (probe with obstacle: difference = -0.10°, [-0.14,-0.05], pd > 99%, 52% in ROPE; probe without obstacle: difference = 0.07°, [0.0,0.15], pd > 97%, 64% in ROPE), whereas there was no statistical difference in Motor Imagery (probe with obstacle: difference = -0.01°, [-0.06,0.04], pd = 64%, 100% in ROPE; probe without obstacle: difference = 0.01°, [-0.06,0.09], pd = 63%, 100% in ROPE). **I-L** Differences in final reach error differences between Execution and Motor Imagery (**IJ**) and No Movement condition (**KL**), respectively. In Experiment 1, the differences in final reach error between trials with and without an obstacle in the prime movement were larger for execution than imagery trials, but only for probe movements with an obstacle (probe with obstacle: difference in final reach error difference = -0.10°, [-0.17,-0.03], pd > 99%, 44% in ROPE; probe without obstacle: difference in final reach error difference = 0.03°, [-0.07,0.14], pd = 74%, 80% in ROPE). In Experiment 2, the differences in final reach error between trials with and without an obstacle in the prime phase were larger for Execution than No Movement, but again only for probe movements with an obstacle (probe with obstacle: difference in final reach error difference = -0.09°, [-0.15,-0.02], pd > 99%, 57% in ROPE; probe without obstacle: difference in final reach error difference = 0.06°, [-0.04,0.16], pd = 89%, 66% in ROPE). Colored areas: posterior distributions. Black dots: median; error bars: 95% highest-density intervals (HDI); grey shaded areas: region of practical equivalence (ROPE).

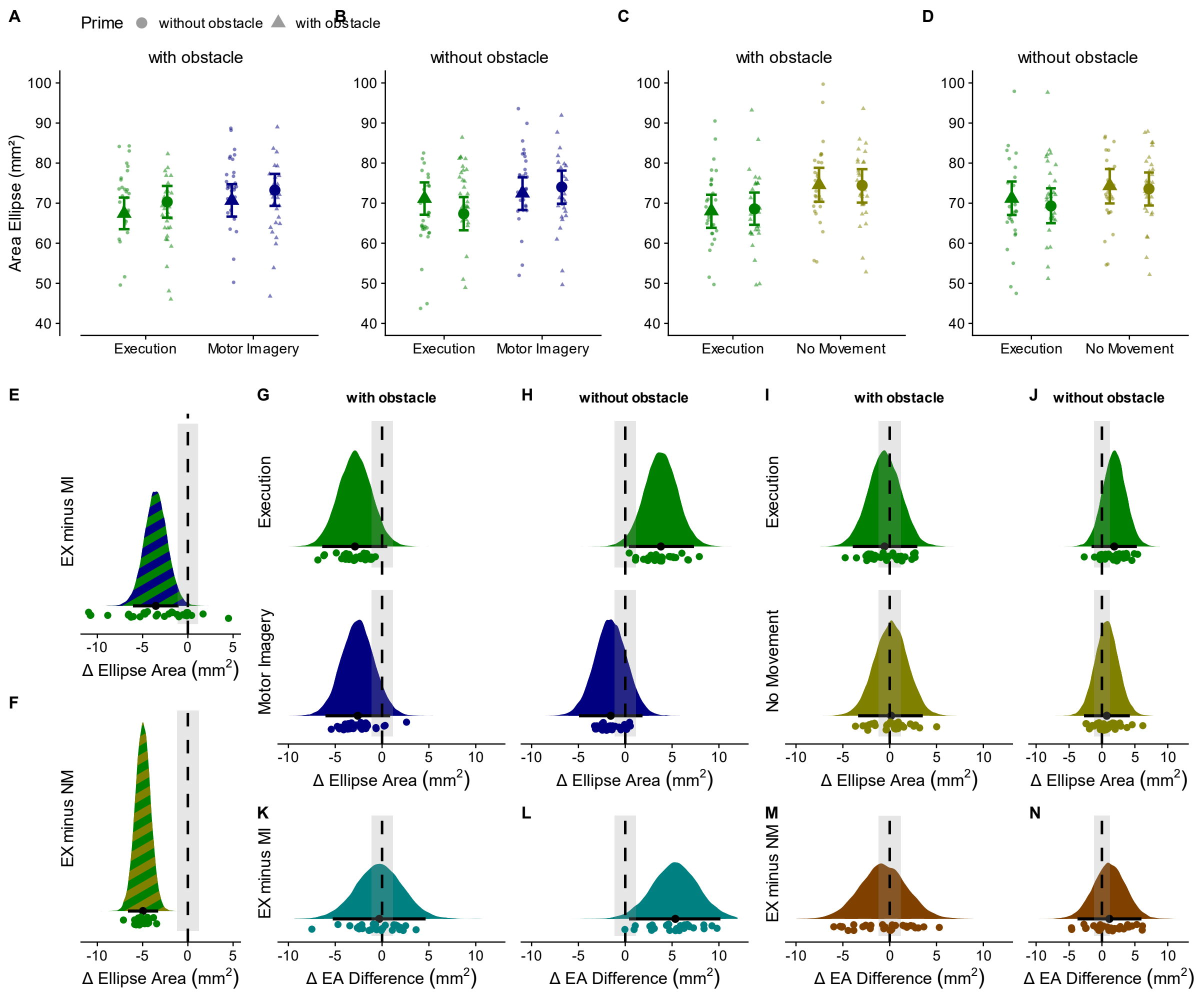


**Figure S4.** Reach endpoint precision of Experiment 1 (Motor Imagery) and Experiment 2 (No Movement). **A-D** Reach endpoint variability for Experiment 1 (AB) and Experiment 2 (CD). **EF** Difference in endpoint variability between Execution and Motor Imagery/ No Movement. Endpoint variability was, on average, smaller following executed compared to imagined or not executed prime movements (Experiment 1: difference = -3.5mm², [-6.1,-1.0], pd > 99%, 3% in ROPE; Experiment 2: difference = -5.0mm², [-6.6,-3.3], pd > 99%, 0% in ROPE). **G-J** Differences in endpoint variability between trials with and without an obstacle in the prime phase for Execution and Motor Imagery/ No Movement with and without an obstacle in the probe movement. In Experiment 1, endpoint variability of execution trials was smaller when prime and probe necessitated similar movements, though statistically reliable only for probe movements without an obstacle (probe with obstacle: difference = -2.9mm², [-6.4,0.6], pd = 95%, 24% in ROPE; probe without obstacle: difference = 3.8mm², [0.4,7.3], pd = 98%, 11% in ROPE), whereas there was no statistical difference in Motor Imagery (probe with obstacle: difference = -2.6mm², [-6.0,0.9], pd = 93%, 29% in ROPE; probe without obstacle: difference = -1.6mm², [-5.0,1.8], pd = 82%, 33% in ROPE). In Experiment 2, no difference was statistically present (Execution probe with obstacle: difference = -0.6mm², [-3.9,3.0], pd = 62%, 35% in ROPE; Execution probe without obstacle: difference = 1.9mm², [-1.6,5.3], pd = 86%, 35% in ROPE; No Movement probe with obstacle: difference = 0.2mm², [-3.4,3.5], pd = 54%, 35% in ROPE; No Movement probe without obstacle: difference = 0.7mm², [-2.7,4.2], pd = 66%, 34% in ROPE). **K-N** Difference in endpoint variability differences between Execution and Motor Imagery **(KL)** and No Movement (**MN**), respectively. In Experiment 1, the differences in endpoint variability between trials with and without an obstacle in the prime phase were larger for execution than imagery trials, but only for probe movements without an obstacle (probe with obstacle: difference in ellipse area difference = -0.3mm², [-5.3,4.7], pd = 55%, 23% in ROPE; probe without obstacle: difference in ellipse area difference = 5.3mm², [0.4,10.2], pd = 98%, 7% in ROPE). In Experiment 2, the differences in endpoint variability between trials with and without an obstacle in the prime phase were statistically equivalent between Execution and No Movement (probe with obstacle: difference in ellipse area difference = -0.6mm², [-5.6,4.3], pd = 61%, 24% in ROPE; probe without obstacle: difference in ellipse area difference = 1.1mm², [-3.7,6.0], pd = 68%, 25% in ROPE). All other details as in Figure S3.

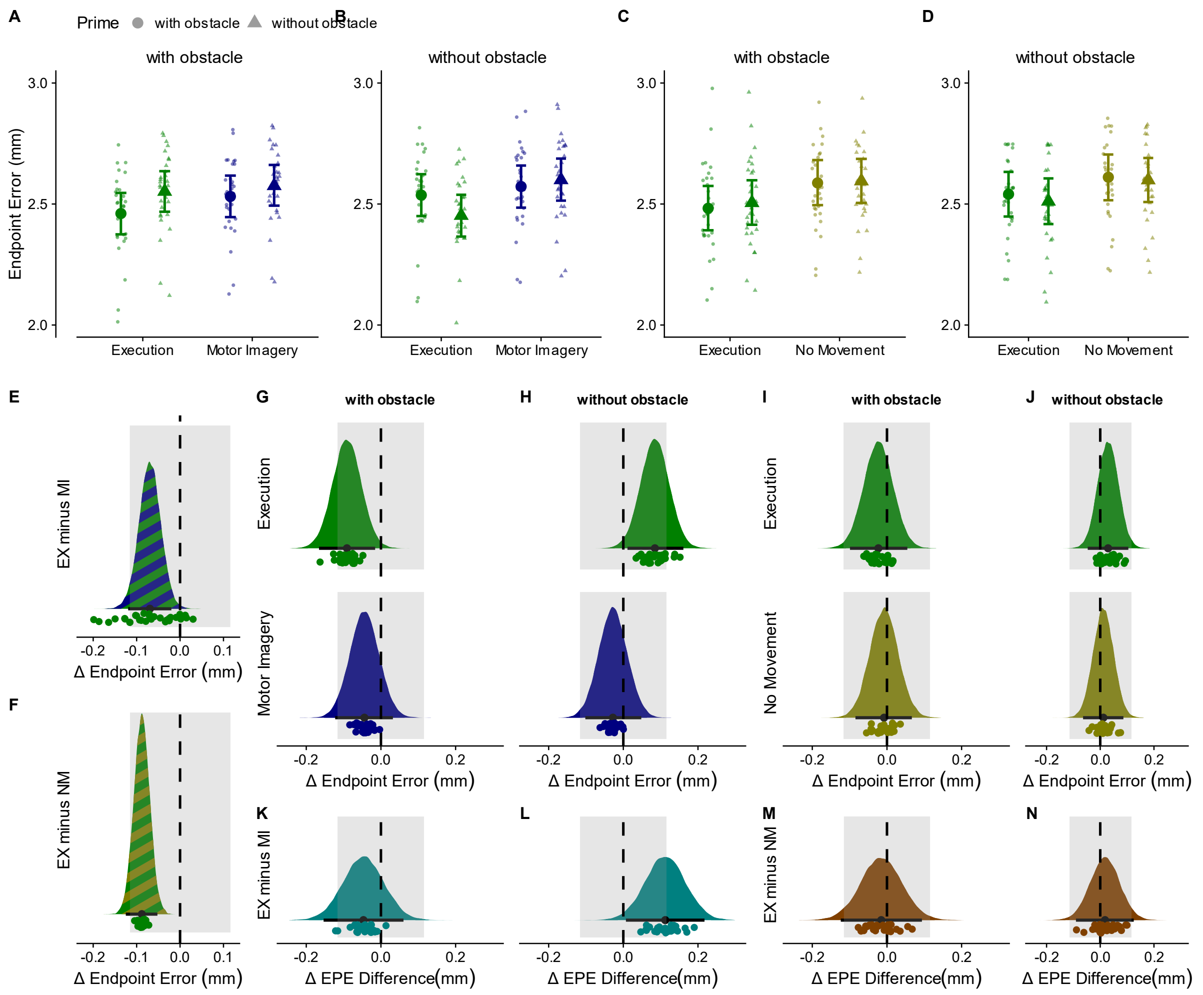


**Figure S5.** Reach endpoint accuracy of Experiment 1 (Motor Imagery) and Experiment 2 (No Movement). **A-D** Reach endpoint error for Experiment 1 (AB) and Experiment 2 (CD). **EF** Endpoint error was, on average, smaller following executed compared to imagined or not executed prime movements. (Experiment 1: difference = -0.07mm, 95% HDI [-0.12,-0.02], pd = 99.60%, 96% of HDI in ROPE; Experiment 2: difference = -0.08mm, 95% HDI [-0.12,-0.05], pd = 100%, 87% of HDI in ROPE). **G-J** Differences in endpoint error between trials with and without an obstacle in the prime phase for Execution and Motor Imagery/ No Movement trials with and without an obstacle in the probe phase. In Experiment 1, endpoint error of execution trials was smaller when prime and probe necessitated similar movements (probe with obstacle: difference = -0.09mm, [-0.17,-0.02], pd > 99%, 67% in ROPE; probe without obstacle: difference = 0.08mm, [0.01,0.16], pd = 99%, 70% in ROPE), whereas there was no statistical difference in imagery trials (probe with obstacle: difference = -0.04mm, [-0.12,0.03], pd = 87%, 96% in ROPE; probe without obstacle: difference = -0.03mm, [-0.10,0.05], pd = 77%, 100% in ROPE). In Experiment 2, no difference was statistically present (Execution probe with obstacle: difference = -0.02mm, [-0.10,0.05], pd = 72%, 100% in ROPE; Execution probe without obstacle: difference = 0.03mm [-0.05,0.10], pd = 78%, 100% in ROPE; No Movement probe with obstacle: difference = -0.01mm, [-0.08,0.07], pd = 58%, 100% in ROPE; No Movement probe without obstacle: difference = 0.01mm, [-0.06,0.08], pd = 62%, 100% in ROPE). **K-N** Difference in endpoint error differences between Execution and Motor Imagery **(KL)** and No Movement trials (**MN**), respectively. In Experiment 1, the differences in endpoint error between trials with and without an obstacle in the prime phase were larger for execution that imagery trials, but only for probe movements without an obstacle (probe with obstacle: difference in endpoint error difference = -0.05mm, [-0.15,0.06], pd = 81%, 83% in ROPE; probe without obstacle: difference in endpoint error difference = 0.11mm, [0.01,0.22], pd = 98%, 52% in ROPE). In Experiment 2, the differences in endpoint error between trials with and without an obstacle in the prime phase were statistically equivalent between Execution and No Movement (probe with obstacle: difference in endpoint error difference = -0.02mm, [-0.12,0.09], pd = 61%, 96% in ROPE; probe without obstacle: difference in endpoint error difference = 0.02mm, [-0.09,0.12], pd = 63%, 96% in ROPE). All other details as in Figure S3.

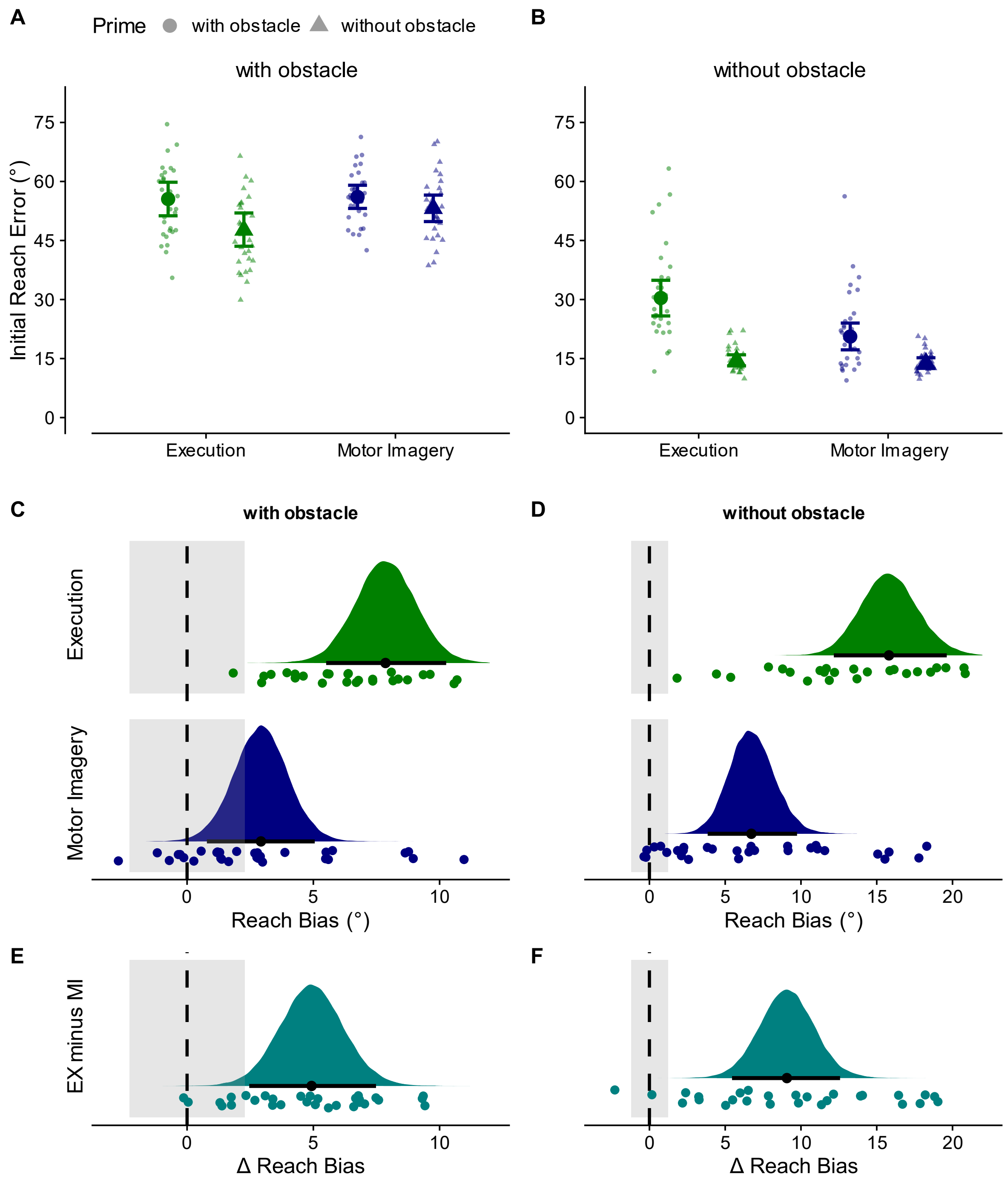


**Figure S6.** Influences of prime movement on feedforward aspects of probe movement in Experiment 3. **AB** Initial reach error. **CD** Reach biases (calculated as difference in initial reach error between trials with and without an obstacle in the prime phase) for execution and motor imagery with and without an obstacle in the probe phase. **EF** Difference in reach biases between Execution and Motor Imagery trials. All other details as in Figure S3.

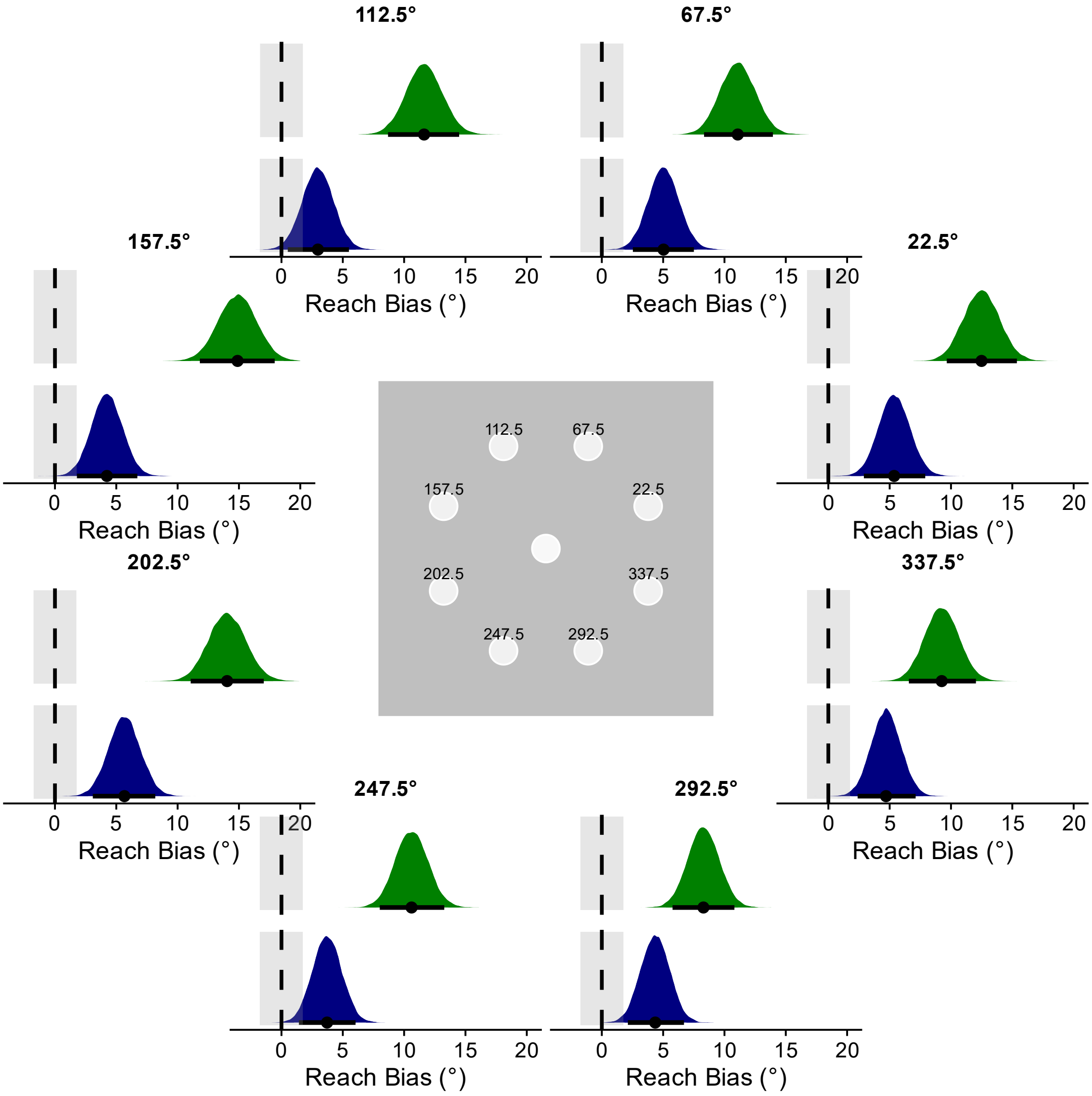


**Figure S7.** Reach biases for the eight targets in Experiment 3 (calculated as difference in initial reach error between trials with and without an obstacle in the prime phase) for Execution and Motor Imagery (averaged across trials with and without an obstacle in probe movements). Biases were present for all target locations. All other details as in Figure S3.

**Table S2:** Reach Bias as a function of trial type and target location in Experiment 3.

| Trial Type | Target Location (°) | Bias | HDI | % in ROPE | pd (%) |
|---|---|---|---|---|---|
| Execution | 22.5 | 12.49 | [9.66,15.37] | 0.0 | 100.0 |
| Motor Imagery | 22.5 | 5.36 | [2.9,7.9] | 0.0 | 100.0 |
| Execution | 67.5 | 11.08 | [8.33,13.95] | 0.0 | 100.0 |
| Motor Imagery | 67.5 | 5.03 | [2.53,7.51] | 0.0 | 100.0 |
| Execution | 112.5 | 11.63 | [8.69,14.48] | 0.0 | 100.0 |
| Motor Imagery | 112.5 | 2.97 | [0.52,5.5] | 24.7 | 99.1 |
| Execution | 157.5 | 14.89 | [11.81,17.92] | 0.0 | 100.0 |
| Motor Imagery | 157.5 | 4.24 | [1.79,6.72] | 0.0 | 100.0 |
| Execution | 202.5 | 14.03 | [11.07,17.02] | 0.0 | 100.0 |
| Motor Imagery | 202.5 | 5.66 | [3.09,8.18] | 0.0 | 100.0 |
| Execution | 247.5 | 10.61 | [8.01,13.27] | 0.0 | 100.0 |
| Motor Imagery | 247.5 | 3.72 | [1.42,6.04] | 7.0 | 99.9 |
| Execution | 292.5 | 8.28 | [5.77,10.81] | 0.0 | 100.0 |
| Motor Imagery | 292.5 | 4.36 | [2.12,6.7] | 0.0 | 100.0 |
| Execution | 337.5 | 9.24 | [6.56,12.03] | 0.0 | 100.0 |
| Motor Imagery | 337.5 | 4.70 | [2.39,7.12] | 0.0 | 100.0 |

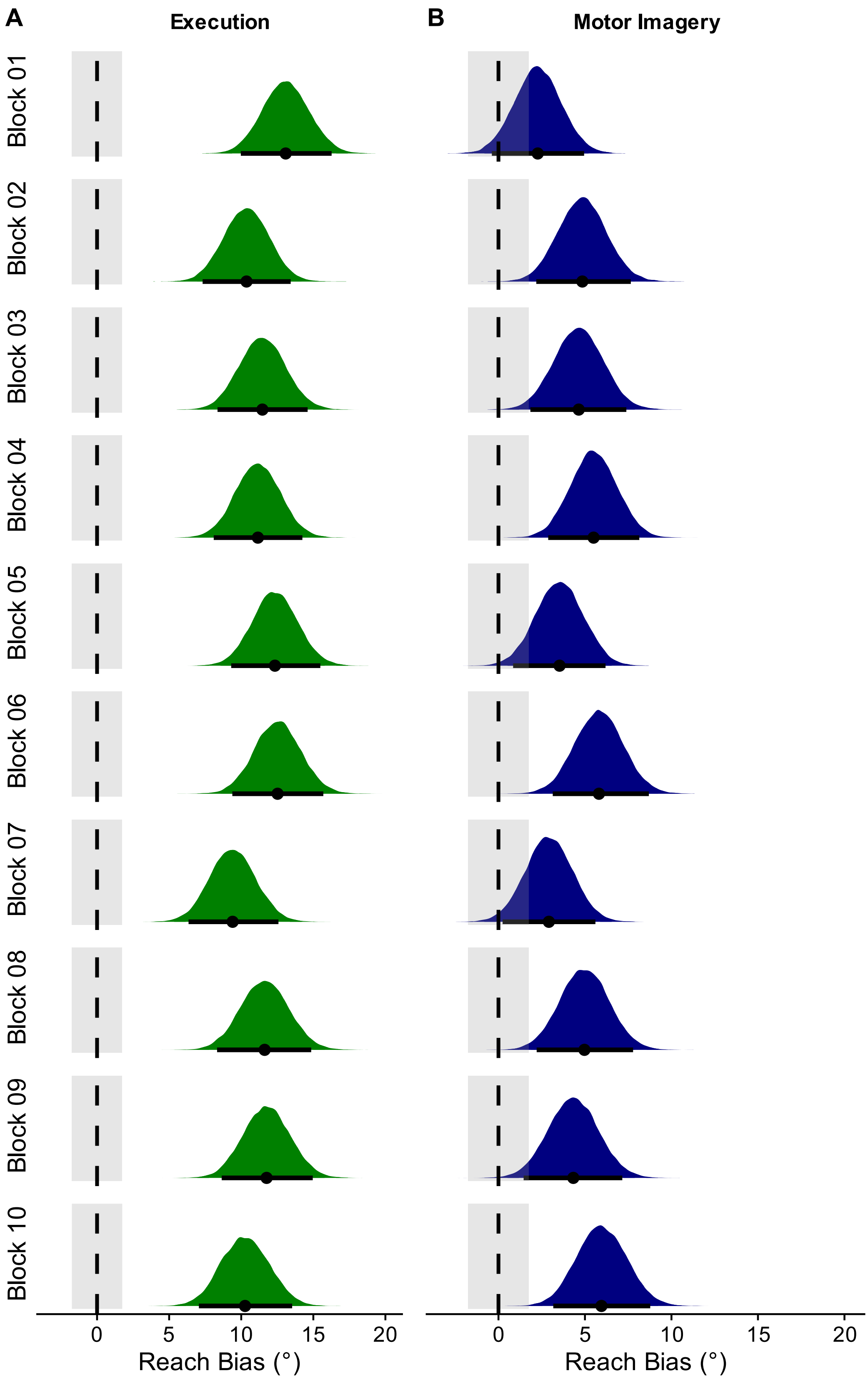


**Figure S8.** Reach biases across the ten blocks in Experiment 3 (calculated as difference in initial reach error between trials with and without an obstacle in the prime phase) for Execution and Motor Imagery (averaged across trials with and without an obstacle in probe movements). All other details as in Figure S3.

**Table S3.** Reach Bias as a function of trial type and block number in Experiment 3.

| Trial Type | Block Number | Bias | HDI | % in ROPE | pd (%) |
|---|---|---|---|---|---|
| Execution | 1 | 13.08 | [9.97,16.27] | 0.00 | 100.0 |
| Motor Imagery | 1 | 2.27 | [-0.38,4.95] | 39.92 | 95.3 |
| Execution | 2 | 10.37 | [7.32,13.43] | 0.00 | 100.0 |
| Motor Imagery | 2 | 4.85 | [2.19,7.65] | 0.00 | 100.0 |
| Execution | 3 | 11.47 | [8.36,14.61] | 0.00 | 100.0 |
| Motor Imagery | 3 | 4.64 | [1.85,7.39] | 0.00 | 100.0 |
| Execution | 4 | 11.15 | [8.09,14.24] | 0.00 | 100.0 |
| Motor Imagery | 4 | 5.50 | [2.88,8.14] | 0.00 | 100.0 |
| Execution | 5 | 12.34 | [9.3,15.49] | 0.00 | 100.0 |
| Motor Imagery | 5 | 3.53 | [0.85,6.19] | 16.84 | 99.5 |
| Execution | 6 | 12.52 | [9.38,15.69] | 0.00 | 100.0 |
| Motor Imagery | 6 | 5.81 | [3.14,8.69] | 0.00 | 100.0 |
| Execution | 7 | 9.40 | [6.35,12.58] | 0.00 | 100.0 |
| Motor Imagery | 7 | 2.91 | [0.24,5.6] | 28.09 | 98.3 |
| Execution | 8 | 11.62 | [8.32,14.85] | 0.00 | 100.0 |
| Motor Imagery | 8 | 4.97 | [2.21,7.78] | 0.00 | 100.0 |
| Execution | 9 | 11.76 | [8.65,14.96] | 0.00 | 100.0 |
| Motor Imagery | 9 | 4.33 | [1.45,7.16] | 5.12 | 99.8 |
| Execution | 10 | 10.26 | [7.06,13.53] | 0.00 | 100.0 |
| Motor Imagery | 10 | 5.95 | [3.17,8.77] | 0.00 | 100.0 |

As in Experiment 1, prime duration was longer in trials with an obstacle for both execution and motor imagery trials (execution: difference = 131 ms, [97,163], pd > 99%, 0% in ROPE; motor imagery: difference = 95 ms, [56,134], pd > 99%, 0% in ROPE; Fig. S9). We again observed no reliable correlation between the reach bias and the Motor Imagery Questionnaire (MIQ-RS) scores (Fig. S10 and Table S4). Finally, as in Experiments 1 and 2, final reach error was influenced when a preceding movement was executed (probe with obstacle: difference = -0.07°, [-0.13,-0.02], pd > 99%, 73% in ROPE; probe without obstacle: difference = 0.07°, [0.00,0.14], pd = 98%, 71% in ROPE; Fig. S11CD green data), but not when it was only imagined (probe with obstacle: difference = -0.01°, [-0.06,0.04], pd = 65%, 100% in ROPE; probe without obstacle: difference = 0.03, [-0.03,0.10], pd = 83%, 99% in ROPE; Fig. S11CD blue data). In addition, reach precision and accuracy was again better following executed compared to imagined trials (difference in ellipse area = -3.0mm$^2$, [-5.8,-0.3], pd = 99%, 13% in ROPE; difference in endpoint error = -0.07mm, [-0.11,-0.02], pd > 99%, 100% in ROPE), though not specifically modulated by prior movement characteristics (all pd < 84%; Fig. S12 & Fig. S13).

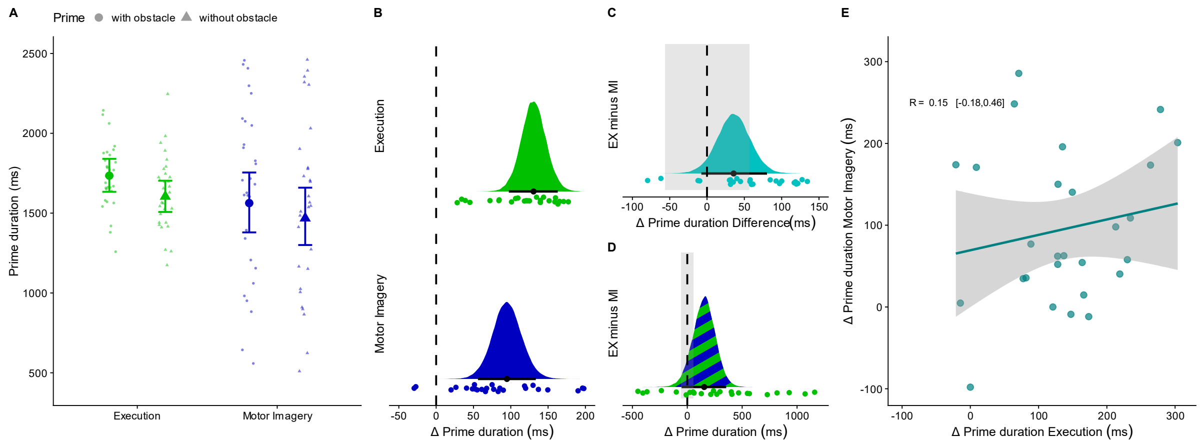


**Figure S9.** Temporal characteristics of prime movements in Experiment 3. **A** Prime duration as a function of trial type (Execution, Motor Imagery) and prime context (with obstacle, without obstacle) in Experiment 3. Large symbols represent the group estimates and error bars the 95% highest-density interval (HDI). Small symbols represent participant estimates. **B-D** Contrasts. Overall, prime duration was longer for execution compared to motor imagery trials (**C**), but for both trial types, prime duration was longer in trials with than without an obstacle (**B**), and this difference was of similar size (**D**). Colored areas show the posterior distributions. Black dots represent the medians and error bars the 95% highest-density intervals (HDI). The gray shaded areas indicate the region of practical equivalence (ROPE). **E** Correlation between difference in prime duration (trials with obstacle – trials without obstacle) in execution and motor imagery trials. The teal line shows a linear fit (±SE, shaded grey area) to the data.

**Table S4.** Correlations between Reach Bias and Motor Imagery Questionnaire Scores (Experiment 3)

| Measure | Trial Type | Probe | r | HDI | pd (%) |
|---|---|---|---|---|---|
| MIQ Score Overall | Execution | without obstacle | 0.04 | [-0.27, 0.36] | 60.27 |
| MIQ Score Overall | Motor Imagery | without obstacle | 0.05 | [-0.27, 0.37] | 61.82 |
| MIQ Score Overall | Execution | with obstacle | 0.01 | [-0.30, 0.33] | 53.20 |
| MIQ Score Overall | Motor Imagery | with obstacle | 0.15 | [-0.17, 0.47] | 80.58 |
| MIQ Score Visual | Execution | without obstacle | 0.07 | [-0.26, 0.37] | 65.70 |
| MIQ Score Visual | Motor Imagery | without obstacle | 0.04 | [-0.33, 0.34] | 59.48 |
| MIQ Score Visual | Execution | with obstacle | -0.01 | [-0.34, 0.33] | 53.17 |
| MIQ Score Visual | Motor Imagery | with obstacle | 0.02 | [-0.31, 0.33] | 54.60 |
| MIQ Score Kinesthetic | Execution | without obstacle | 0.02 | [-0.30, 0.34] | 54.97 |
| MIQ Score Kinesthetic | Motor Imagery | without obstacle | 0.03 | [-0.28, 0.38] | 58.03 |
| MIQ Score Kinesthetic | Execution | with obstacle | 0.04 | [-0.30, 0.34] | 58.53 |
| MIQ Score Kinesthetic | Motor Imagery | with obstacle | 0.23 | [-0.10, 0.53] | 90.12 |
| Ease of Imagery | Execution | without obstacle | 0.08 | [-0.23, 0.40] | 68.27 |
| Ease of Imagery | Motor Imagery | without obstacle | 0.14 | [-0.20, 0.44] | 79.12 |
| Ease of Imagery | Execution | with obstacle | -0.03 | [-0.33, 0.29] | 56.62 |
| Ease of Imagery | Motor Imagery | with obstacle | 0.15 | [-0.18, 0.47] | 80.10 |
| Count of Imagery | Execution | without obstacle | 0.01 | [-0.30, 0.32] | 52.35 |
| Count of Imagery | Motor Imagery | without obstacle | 0.19 | [-0.15, 0.47] | 86.22 |
| Count of Imagery | Execution | with obstacle | 0.05 | [-0.27, 0.36] | 61.35 |
| Count of Imagery | Motor Imagery | with obstacle | 0.15 | [-0.18, 0.45] | 80.80 |

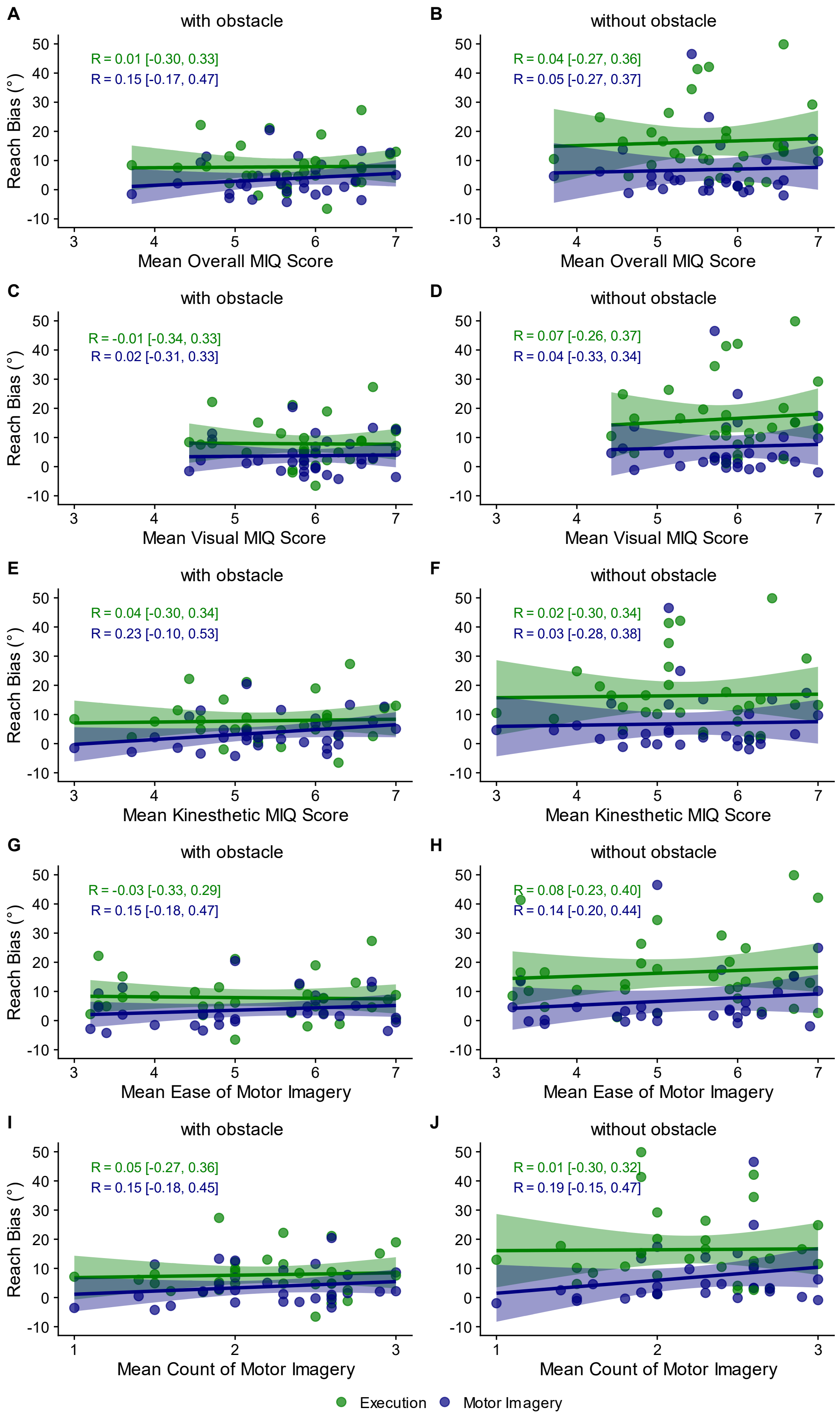


**Figure S10.** Experiment 3. Correlation between Reach Bias and **AB** Mean Overall MIQ Score, **CD** Mean Visual MIQ Score, **EF** Mean Kinesthetic MIQ Score, **GH** Mean Ease of Motor Imagery Score, and **IJ** Mean Count of Motor Imagery Score. Solid lines show linear fits (±SE, shaded area) to the data.

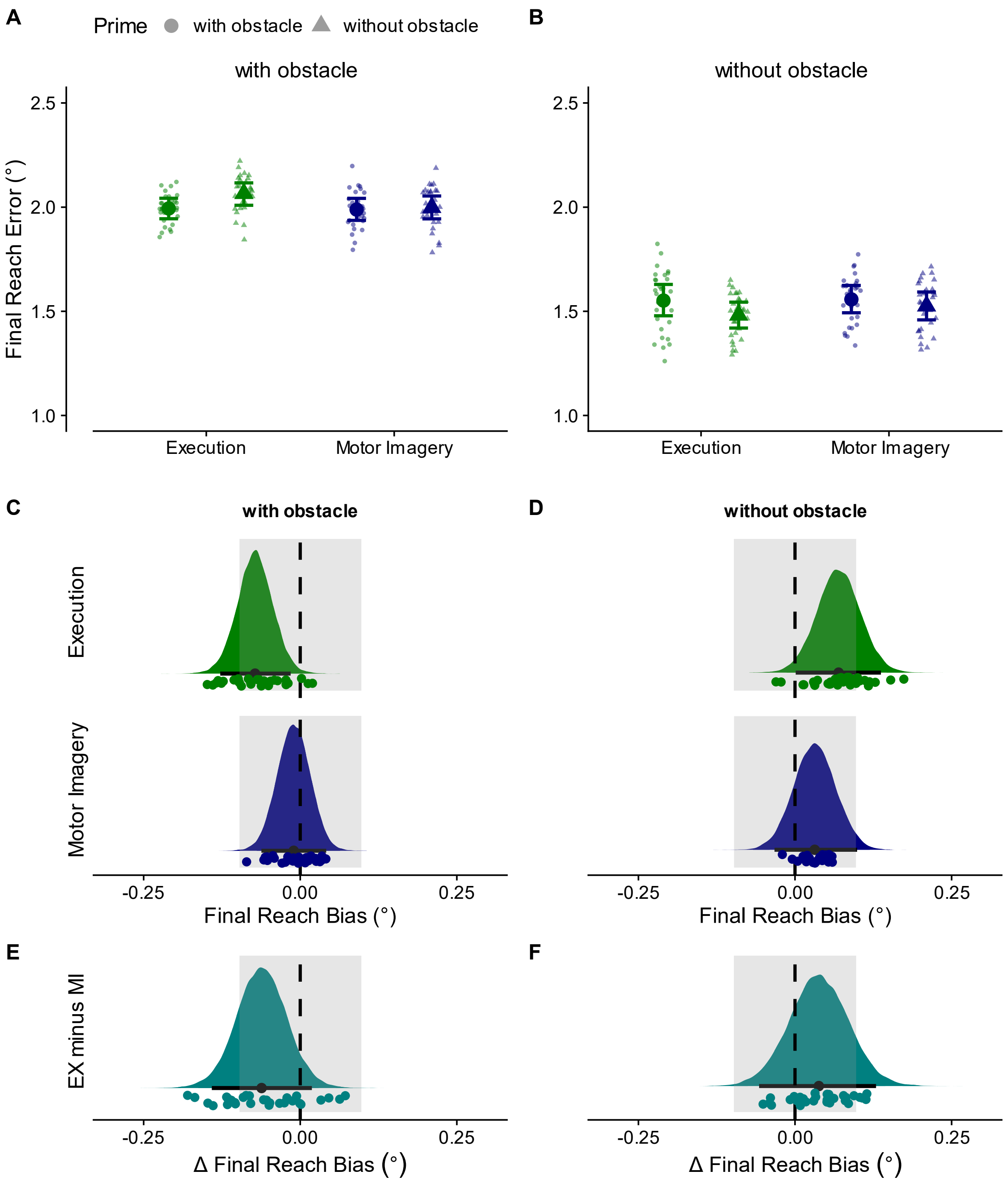


**Figure S11.** Influences of prior movement on feedback aspects in Experiment 3. **AB** Final reach error. **CD** Differences in final reach error between trials with and without an obstacle in the prime phase for Execution and Motor Imagery with and without an obstacle in probe movements. **EF** Difference in final reach error differences between Execution and Motor Imagery. All other details as in Figure S3.

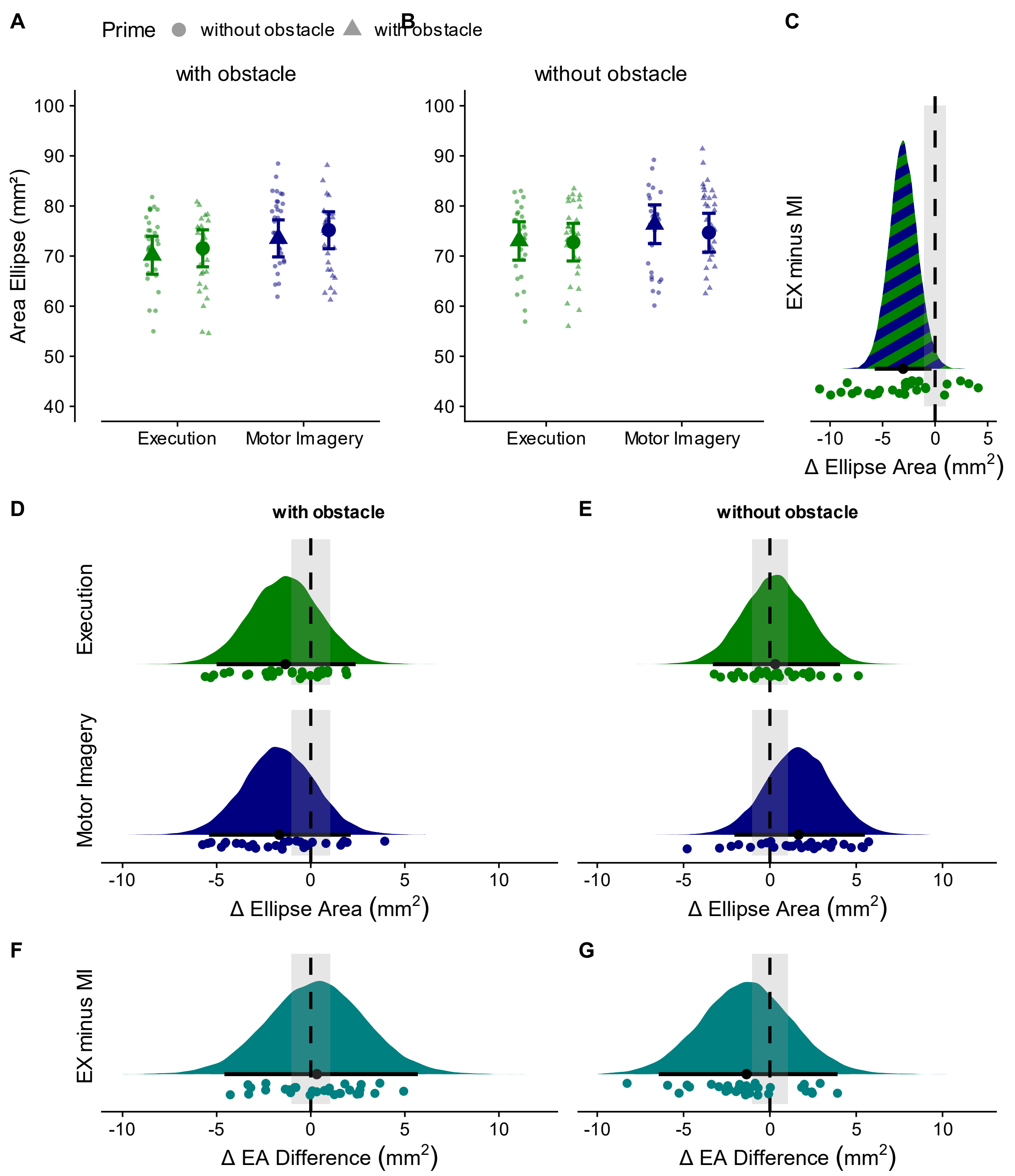


**Figure S12.** Reach precision in Experiment 3. **AB** Reach endpoint precision calculated as the area of the 95% confidence ellipse (in mm²) for reach endpoints 150ms after the cursor entered the target. Endpoint precision was, on average, higher following executed compared to imagined prime movements (difference in ellipse area = -3.0mm², 95% HDI [-5.8,-0.3], pd = 98.50%, 13% of HDI in ROPE). **DE** Differences in ellipse areas between trials with and without an obstacle in the prime phase for execution and motor imagery trials with and without an obstacle in the probe phase. Ellipse areas did statistically not differ (Execution probe with obstacle: difference = -1.4mm², [-5.0,2.4], pd = 77%, 28% in ROPE; Execution probe without obstacle: difference = 0.3mm², [-3.3,4.1], pd = 57%, 28% in ROPE; Motor Imagery probe with obstacle: difference = -1.7mm², [-5.4,2.1], pd = 81%, 27% in ROPE; Motor Imagery probe without obstacle: difference = 1.7mm², [-2.1,5.5], pd = 81%, 27% in ROPE). **FG** Difference in ellipse area differences between Execution and Motor Imagery. The differences were statistically equivalent (probe with obstacle: difference in area difference = 0.3mm², [-4.6,5.7], pd = 55%, 20% in ROPE; probe without obstacle: difference in area difference = -1.4mm², [-6.4,3.9], pd = 70%, 20% in ROPE). All other details as in Figure S3.

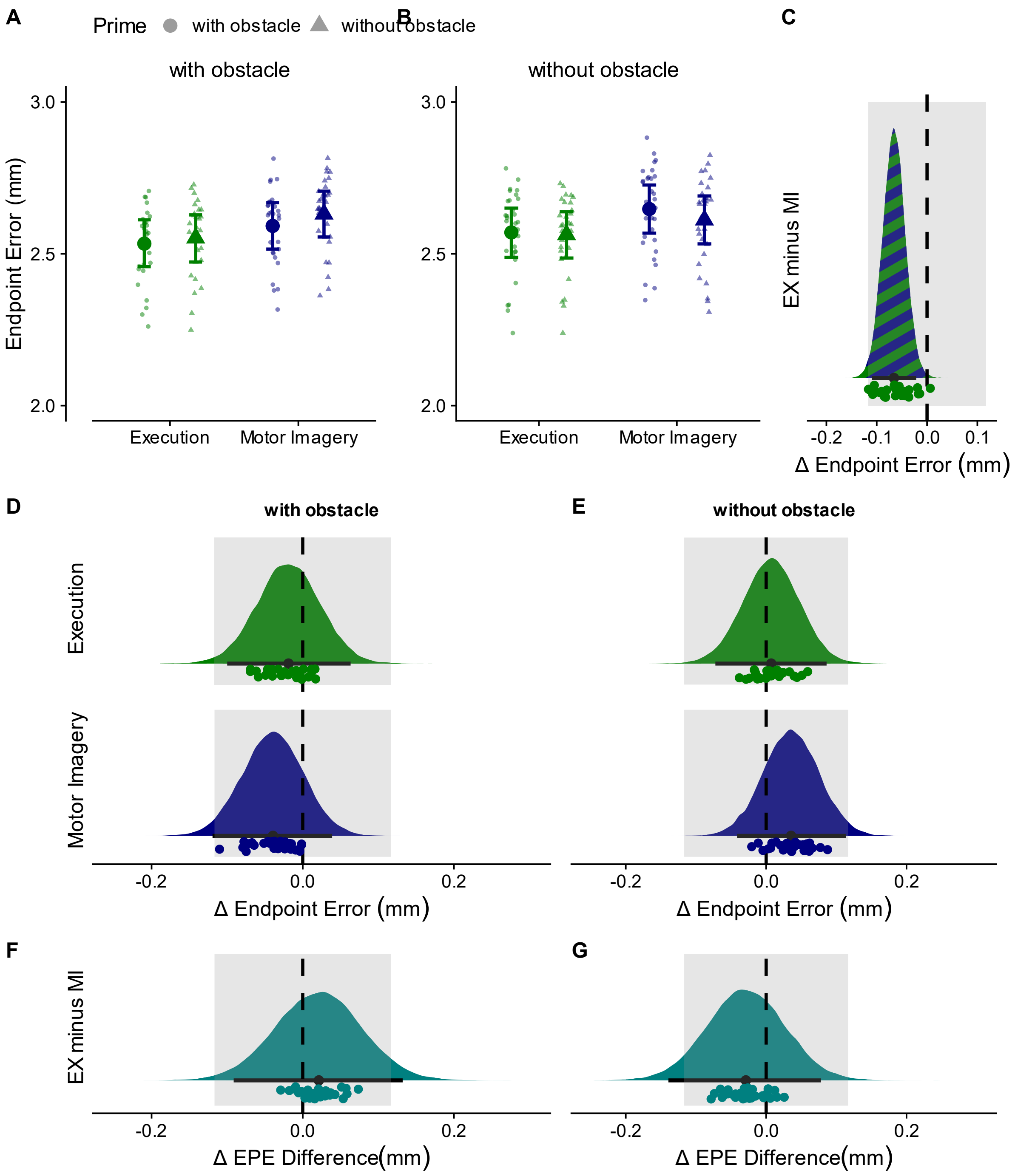


**Figure S13.** Reach accuracy in Experiment 3. **AB** Reach endpoint errors calculated as Euclidean distance of the cursor position 150ms after it entered the target and the target center. Endpoint errors were, on average, smaller following executed compared to imagined prime movements (difference in endpoint error = -0.07mm, [-0.11,-0.02], pd > 99%, 100% in ROPE). **DE** Differences in endpoint errors between trials with and without an obstacle in the prime phase for execution and motor imagery trials with and without an obstacle in the probe phase. Endpoint errors did statistically not differ (Execution probe with obstacle: difference = -0.02mm, [-0.10,0.06], pd = 68%, 100% in ROPE; Execution probe without obstacle: difference = 0.01mm, [-0.07,0.09], pd = 57%, 100% in ROPE; Motor Imagery probe with obstacle: difference = -0.04mm, [-0.12,0.04], pd = 84%, 98% in ROPE; Motor Imagery probe without obstacle: difference = 0.04mm, [-0.04,0.11], pd = 82%, 100% in ROPE). **FG** Difference in endpoint differences between Execution and Motor Imagery. The differences were statistically equivalent (probe with obstacle: difference in endpoint difference = 0.02mm, [-0.09,0.13], pd = 64%, 93% in ROPE; probe without obstacle: difference in endpoint difference = -0.03mm, HDI [-0.14,0.08], pd = 70%, 90% in ROPE). All other details as in Figure S3.

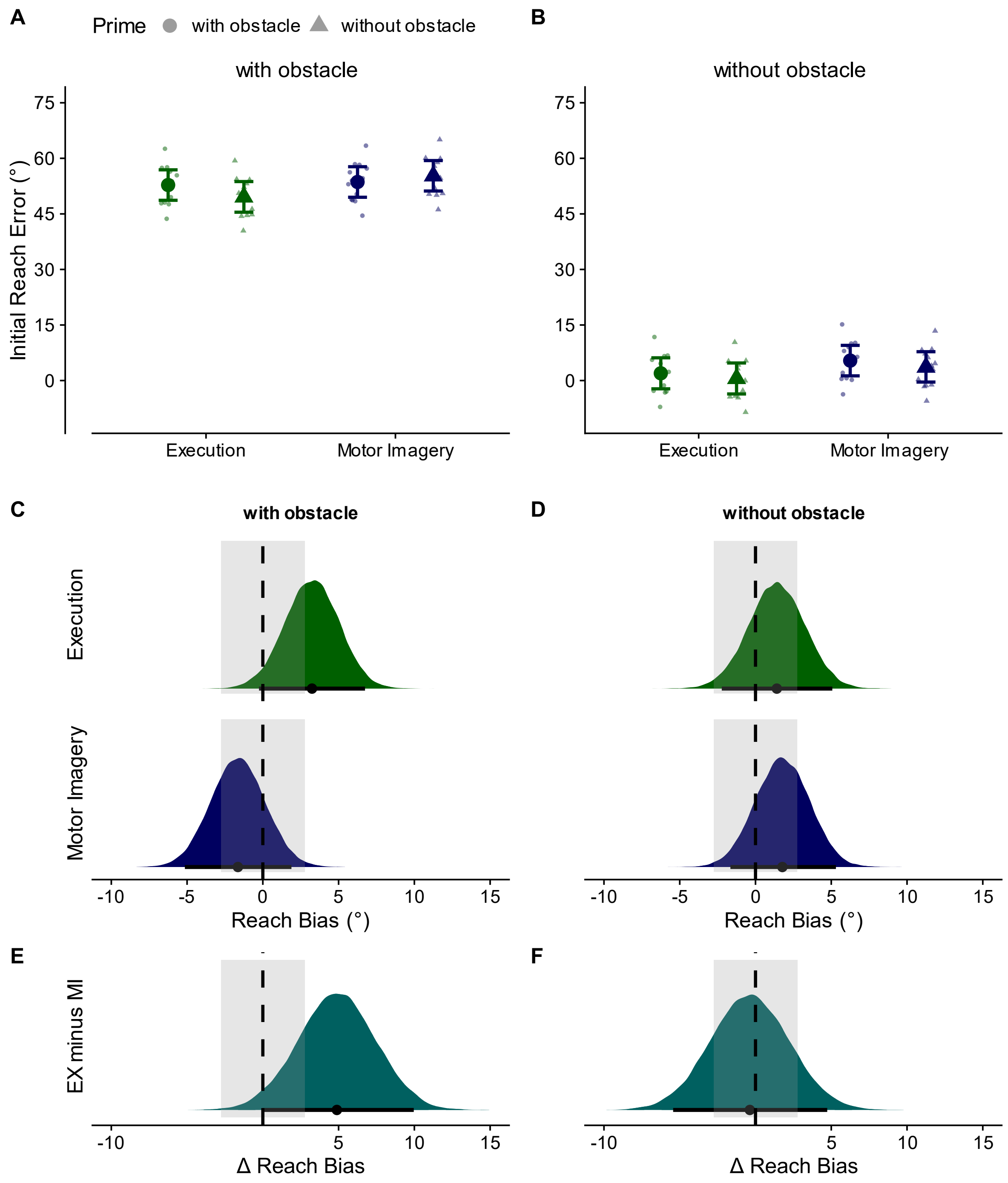


**Figure S14.** Influences of prior movement on feedforward aspects in Roberts et al. (2025). **AB** Initial reach error. **CD** Reach biases (calculated as difference in initial reach error between trials with and without an obstacle in the prime phase) for execution and motor imagery with and without an obstacle in the probe phase. **EF** Difference in reach biases between Execution and Motor Imagery trials. All other details as in Figure S3.
